# Disordered protein COSA-2 maintains crossover-specific repair compartments to ensure meiotic crossover maturation

**DOI:** 10.64898/2026.05.13.725012

**Authors:** Celja J. Uebel, Dahlia Y. Deng, Yumi Kim, Anne M Villeneuve

## Abstract

Faithful genome inheritance during meiosis relies on crossover repair of double-strand DNA breaks (DSBs) to connect homologous chromosomes and direct their proper segregation. The formation of crossover-specific recombination intermediates and accumulation of pro-crossover factors occurs at an extremely limited subset of DSB sites, necessitating that the subset of recombination sites designated to become crossovers reliably mature into crossovers. Here we identify *C. elegans* disordered protein COSA-2 as crucial for meiotic crossover maturation. COSA-2 abruptly concentrates at crossover intermediates in late pachytene nuclei, where it colocalizes and associates with other pro-crossover factors. COSA-2 is dispensable for early loading of crossover factors and for crossover designation, but is required for maintenance of pro-crossover factors at crossover-designated sites and for focal enrichment of factors initially distributed throughout the synaptonemal complex. We define a COSA-2 execution point during late pachytene wherein crossover intermediates transition from a vulnerable state (in which they require COSA-2 to avoid being dismantled) to a state where COSA-2 and local crossover-factor enrichment are no longer required to connect homologs. We propose that COSA-2 scaffolds privileged DNA repair compartments that promote crossover-factor accumulation and protect crossover intermediates until completion of repair, thereby ensuring that crossover-designated sites reliably mature into crossovers.

## Introduction

Crossover (CO) repair of DNA double-strand breaks (DSBs) creates temporary physical connections between homologous chromosome pairs that direct their accurate segregation in the first meiotic division. Failure to form a CO-based connection between a pair of homologs can result in chromosomal aneuploidy, a leading cause of miscarriage and congenital disorders in humans (reviewed in Gruhn and Hoffmann 2022). Thus, mechanisms that promote and ensure the formation of interhomolog CO recombination events are central for reproductive success.

The process of meiotic recombination is initiated during meiotic prophase by the programmed formation of DSBs by the topoisomerase-like Spo11 protein (Keeney et al. 1997; reviewed in Lam and Keeney 2014). DSBs are resected to yield overhanging 3’ end strands, which can invade a homologous template to yield a repair intermediate containing a displacement loop, or “D-loop” (Hunter and Kleckner 2001; Hunter 2015). Following synthesis primed by the invading 3’ end, the vast majority of D-loop intermediates are disassembled and repair is completed to yield non-COs (reviewed in Sanchez et al. 2021a). This non-CO synthesis-dependent strand annealing (SDSA) repair mechanism restores genomic integrity but does not connect homologous chromosomes in a manner that will persist beyond prophase. A crucial but limited subset of D-loop intermediates undergo “second end capture”, in which the other resected end of the DSB anneals to the displaced strand of the joint molecule to form a double Holiday Junction (dHJ)(Schwacha and Kleckner 1995; Hunter and Kleckner 2001). These dHJs are considered “CO-specific intermediates”(Allers and Lichten 2001; Börner et al. 2004), as subsequent resolution of dHJs by structure-selective endonucleases can yield CO recombination products(reviewed in Hunter 2015; Sanchez et al. 2021a). Importantly, dHJs are a necessary precursor to CO resolution, but without appropriate stabilization, they can be prematurely or inappropriately dissolved or dismantled (e.g. by helicase/topoisomerase complexes and/or nonspecific resolvases) to yield non-CO repair products that fail to connect homologs (reviewed in Sanchez et al. 2021a).

Despite the importance of CO-based connections for directing homolog segregation, CO formation is highly constrained in most organisms. The number of meiotic DSBs generated far exceeds the eventual number of COs produced, and CO-specific intermediates form at an extremely limited subset of DSB repair sites, often one per chromosome pair (reviewed in Hunter 2015; Gray and Cohen 2016). Thus, to ensure that sufficient COs are formed to direct accurate chromosome segregation, it is crucial that the DSB repair intermediates that are selected to become CO intermediates reliably mature into CO products. However, the mechanisms that guarantee reliable maturation of CO intermediates into CO products are poorly understood.

Recent work in the model organism *C. elegans*, which relies on a single CO per chromosome pair (Hillers and Villeneuve 2003), has begun to illuminate how CO-specific DNA repair intermediates may be stabilized and protected to promote CO-biased resolution, providing insight into the mechanisms that ensure robustness to the process of meiotic CO formation. First, SIM images of DNA repair and recombination factors localized at CO-designated recombination sites have revealed a unique spatial organization in which the conserved MutSγ meiotic sliding clamp complexes and BLM helicases occupy distinct domains of the underlying DNA intermediates (Jagut et al. 2016; Woglar and Villeneuve 2018). At these sites, doublets of HIM-6/BLM foci were interpreted to represent cohorts of helicase complexes localizing at or near the two junctions of a dHJ intermediate, with orthogonally-oriented doublets of MSH-5 foci representing two populations of MutSγ complexes embracing the two DNA duplexes spanning between the junctions, where they could act as a roadblock to inhibit dHJ dissolution and thereby favor CO resolution (Woglar and Villeneuve 2018). Additional observations suggest that CO sites represent spatially-segregated compartments within the central region of the synaptonemal complex (SC), the highly-ordered yet dynamic structure that forms at the interface between aligned meiotic chromosomes. SIM images have revealed SC central region proteins encasing assemblages of CO factors accumulated at CO sites (Woglar and Villeneuve 2018), a conformation proposed to enable distinct repair outcomes at CO versus non-CO sites by favoring local enrichment of factors promoting CO maturation inside the “SC bubbles” and/or by protecting CO-specific intermediates from non-CO repair activities acting outside the “SC bubbles”. Further, there is substantial evidence for a self-reinforcing positive feedback loop operating at CO-designated sites to recruit and/or retain pro-CO factors, as initially proposed by Yokoo et al. (2012). This positive feedback mechanism involves scaffold-like properties and phosphorylation of intrinsically-disordered C-terminal tails of MSH-5 and the RING-finger proteins ZHP-3/ZHP-4 by the Cyclin-dependent kinase CDK-2 in complex with its cyclin-like partner COSA-1, as well as physical interactions of COSA-1 with MSH-5 and with ZHP-3, that together confer robustness to the process of CO formation (Haversat et al. 2022; Yang et al. 2024; Zhang et al. 2025). Finally, it was recently discovered that COSA-1 interacts directly with the SLX-4 protein that scaffolds HJ resolvases, providing a means to couple designation of CO sites with the factors that execute their eventual resolution (Liu et al. 2026).

Here we introduce a new component of the *C. elegans* CO machinery, COSA-2 (<u>C</u>ross<u>o</u>ver-<u>s</u>ite-<u>a</u>ssociated-2) that we identified in an updated version of our “Green eggs & Him” genetic screen for meiotic factors (Kelly et al. 2000). COSA-2 is an intrinsically disordered protein that localizes to CO sites and plays a unique role in promoting and ensuring the maturation of CO intermediates into COs. Our genetic and cytological analyses reveal that COSA-2 is dispensable for early loading of CO factors at DSB repair sites and for the designation of a subset of these sites to become COs. Instead, COSA-2 is specifically required to retain CO factors at CO-designated sites and to promote local enrichment at CO sites of factors initially distributed throughout the SC, ultimately leading to the formation of CO products that connect homologs and direct their segregation. Collectively, our data support a model in which COSA-2 serves as “molecular glue” and/or “insulation” to scaffold and actively maintain privileged DNA repair compartments in which CO factors accumulate and protect CO intermediates from being disassembled and repaired as non-COs, thereby ensuring that each meiotic DSB repair intermediate selected to initiate CO formation reliably matures as a functional CO product.

## RESULTS

### The unstructured protein COSA-2 is required for CO formation

We identified COSA-2 (K01C8.5, previously known as GEI-14) as a new component of the meiotic machinery based on isolation of two *cosa-2* mutant alleles in an updated version of a genetic screen for mutations causing missegregation of sex chromosomes (Kelly et al. 2000). We created a *cosa-2* null allele, *cosa-2(me169),* by inserting a STOP-IN cassette (Wang et al. 2018) to introduce stop codons in all reading frames (**Figure 1A**); unless otherwise indicated, *cosa-2*(*me169*) was used for all experiments.

**Figure 1.**
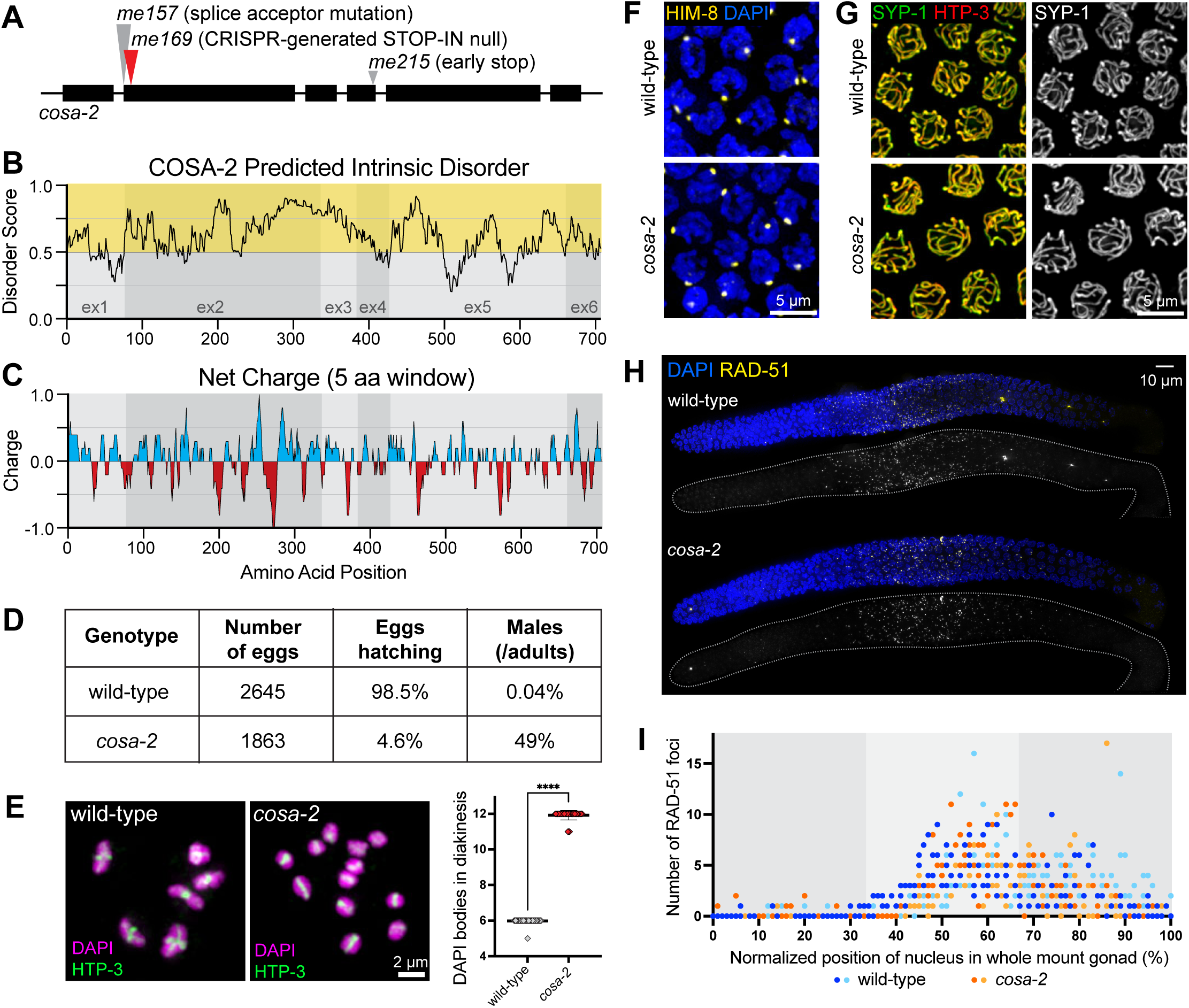
Disordered protein COSA-2 is required for meiotic crossover formation. (A) Schematic of the *cosa-2* gene, indicating positions of the engineered null mutation used throughout this study (red arrow) and point mutations initially identified by genetic screening (gray arrows). (B) Intrinsic disorder values (IUPred3 long disorder parameters (Erdos et al. 2021)) plotted as a function of amino acid position. Values above 0.5 (yellow zone) are categorized as intrinsically disordered and comprise 77.5% of the COSA-2 protein. Light and dark gray regions denote exon boundaries. (C) Net charge of a sliding window of 5 amino acid residues plotted as a function of amino acid position (Holehouse et al. 2017). (D) Quantification of decreased hatching rate and increased male frequency among progeny of *cosa-2* mutant worms. (E) Left, images of full karyotypes of chromosomes in individual diakinesis oocytes, stained for DNA (DAPI) and chromosome axis protein HTP-3. Six DAPI-stained bodies in the wild-type nucleus correspond to 6 bivalents (chromosome pairs connected by chiasmata), whereas 12 DAPI-stained bodies (univalents) are detected in the *cosa-2* mutant nucleus. Right, quantification of DAPI-stained bodies in diakinesis nuclei from wild-type oocytes (n = 53) and *cosa-2* oocytes (n = 52), showing mean ± SD; ***Mann-Whitney p-value < 0.0001. (F) Immunofluorescence images of paring center binding protein HIM-8 in early pachytene nuclei. A single HIM-8 focus per nucleus indicates appropriate pairing of X-chromosome pairing centers in *cosa-2* mutants. (G) Immunofluorescence images of late pachytene nuclei, showing contiguous tracks of SC central region protein SYP-1 colocalizing with chromosome axis marker HTP-3, indicating successful synapsis in the *cosa-2* mutant. (H) Whole-mount gonads immunostained for RAD-51, a DNA strand exchange protein that marks early DSBR intermediates. (I). Quantification of the distributions of RAD-51 foci in wild-type and *cosa-2* mutant germ lines. Each data point represents an individual nucleus, with its normalized position calculated as the percentage of the length between distal tip of the germ line (0%) and the end of pachytene zone (100%); light and dark colors for each genotype represent two different gonads. RAD-51 foci appeared with normal timing and accumulated at normal levels in the middle third of the gonad in the *cosa-2* mutant (Mann-Whitney p=0.452, n.s.); RAD-51 foci declined in abundance in the last third of the gonads in both control and the *cosa-2* mutant, but were modestly reduced in the *cosa-2* mutant relative to control (p=0.007).

*cosa-2* encodes a 707 amino acid protein containing multiple disordered and low complexity domains distributed throughout its entire length, with 77% of the protein predicted to be intrinsically disordered (**Figure 1B, Supplemental Figure S1**). The protein is characterized by alternating peaks of positive and negative charges, with a striking positive-negative-positive peak pattern beginning at amino acid position 245 in which the negative peak contains a string of 7 consecutive glutamic acid residues (**Figure 1C**). COSA-2 orthologs within the *Caenorhabditis* genus exhibit relatively low levels of overall sequence conservation but share similar patterns of predicted disorder, including a broad peak of intrinsic disorder spanning amino acid positions 200-350 (**Supplemental Figure S1**), suggesting that disorder and charge patterning may confer protein functionality.

*cosa-2* mutants exhibit phenotypes characteristic of mutants defective in meiotic recombination. Self-fertilizing *cosa-2* hermaphrodites (XX) produce a high frequency of inviable embryos (95.3%, n = 1863), and nearly half of their surviving progeny are XO male (49%, n = 85), reflecting missegregation of both autosomes and sex chromosomes (**Figure 1D**). Further, whereas late diakinesis-stage oocytes in wild-type animals contain 6 compact bivalents (chromosome pairs connected via crossover-based connections known as chiasmata), diakinesis oocytes in *cosa-2* animals contain 12 univalents (**Figure 1E**), indicating an absence of chiasmata at the end of meiotic prophase.

### COSA-2 is a component of the meiotic crossover machinery

Several lines of evidence together implicate COSA-2 as a component of meiotic CO machinery. First, immunolocalization experiments indicate that *cosa-2* mutants are proficient for pairing and synapsis between homologous chromosomes. As in wild type, X-chromosome pairing center protein HIM-8 localizes to a single bright focus in pachytene nuclei in *cosa-2* mutants, indicating successful X chromosome pairing (**Figure 1F**). Further, *cosa-2* mutants display contiguous linear stretches of SYP-1, an SC central region protein, colocalizing with meiotic chromosome axis component HTP-3, indicating successful synapsis (**Figure 1G**). Second, immunostaining for DNA strand exchange protein RAD-51, which marks early DSB-dependent recombination intermediates, showed that RAD-51 foci appear with normal timing and reach normal levels in *cosa-2* mutant germ lines before declining in number (**Figure 1H,I**). These data indicate that *cosa-2* mutants are competent to generate the DSBs that serve as the initiating events of meiotic recombination, but are not successful at completing DSB repair in a manner that leads to interhomolog COs.

Third, COSA-2 exhibits a localization pattern characteristic of proteins that concentrate at CO-designated recombination sites. COSA-2::3XFLAG expressed from the endogenous *cosa-2* locus (*cosa-2::3xFLAG)* is detected as foci in nuclei beginning at the onset of the late pachytene stage of meiotic prophase, localizing to a single focus per chromosome pair (6 total foci per nucleus), which corresponds to the number of CO-designated sites present at this stage (**Figure 2Ai, 2Aii**). As nuclei progress into the diplotene stage and the SC central region proteins become asymmetrically redistributed around the presumptive crossover site (Nabeshima et al. 2005), the single COSA-2 focus is located at the boundary between the “short arm” (here marked by SYP-1 accumulation) and long arm (where SYP-1 is depleted) of the remodeling bivalents (**Figure 2Aiii**). As nuclei progress into diakinesis, the final stage of meiotic prophase, COSA-2 foci are found at the intersection of the cruciform HTP-3 signal, a site corresponding to chromosomal remodeling around chiasmata, the CO-based linkage between the homologs (**Figure 2Aiv**). This localization of COSA-2 from the late pachytene stage onward closely parallels that of COSA-1, a cyclin-like protein that partners with the cyclin dependent kinase CDK-2 and localizes in bright foci at sites of eventual COs (**Figure 2B**) (Yokoo et al. 2012; Haversat et al. 2022). Diplotene COSA-2 foci similarly colocalize with the focal accumulation of RING finger protein ZHP-3 that occurs at the short-arm/long-arm boundary (**Supplemental Figure S2**) (Jantsch et al. 2004; Bhalla et al. 2008).

**Figure 2.**
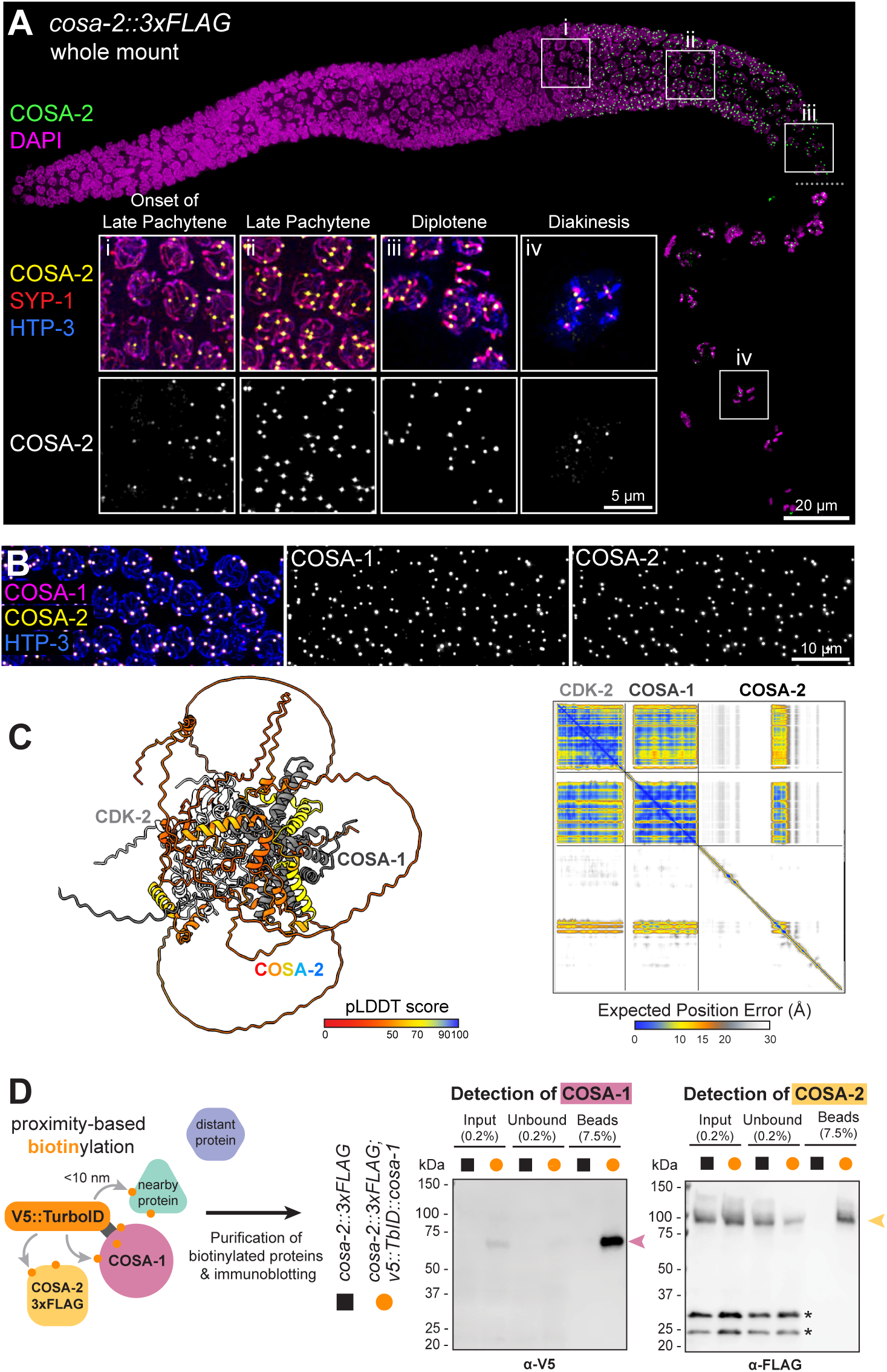
COSA-2 localizes to CO sites and exhibits close molecular proximity to COSA-1. (A) Immunostained whole mount gonad showing the localization of COSA-2::3xFLAG as chromosome-associated foci from late pachytene onset through the diakinesis stage. Boxes indicate the position of insets i-iv.; a small dashed line denotes where a different DAPI contrast adjustment was used for ideal diakinesis DNA visualization. Insets show i-ii) onset of COSA-2::3xFLAG foci as nuclei transition from mid-pachytene to late pachytene, iii) localization of the COSA-2 focus at the boundary between SYP-enriched and SYP-depleted domains at the diplotene stage, and iv) persistence of COSA-2::3xFLAG foci into diakinesis, where a focus colocalizes with the chiasma site on each chromosome pair. (B) Nuclear spread preparation of *cosa-2::3xFLAG; OLLAS::cosa-1* gonads show colocalization of OLLAS::COSA-1 and COSA-2::3XFLAG. (C) Left: AlphaFold3 model of the CDK-2–COSA-1–COSA-2 complex, with COSA-2 colored by predicted Local Distance Difference Test (pLDDT) score and CDK-2 and COSA-1 shown in gray. Right: The Predicted Aligned Error (PAE) matrix indicates high-confidence interactions between COSA-1 and COSA-2 in this model. (D) Schematic of TurboID proximity labeling, wherein TurboID fused to COSA-1 biotinylates proteins within 10 nm. Immunoblots detecting enrichment of V5::TurboID::COSA-1 (expected MW: 76 kDa) and COSA-2::3xFLAG (expected MW: 85 kDa) following affinity purification of biotinylated proteins from worms co-expressing the two proteins. Asterisk indicates non-specific band.

Given the colocalization of COSA-2 with COSA-1 at CO sites, we used AlphaFold3 to predict potential direct interaction between these proteins (Abramson et al. 2024). The prediction yielded a high-confidence model in which COSA-2 binds COSA-1 in a ternary complex with CDK-2 (**Figure 2C, Supplemental Figure 3**). Although predicted Local Distance Difference Test (pLDDT) scores are low across the disordered regions of COSA-2, the two alpha-helices that contact the hydrophobic groove of COSA-1 are predicted with high confidence (pLDDT>70). To test whether COSA-2 and COSA-1 are in close proximity in vivo, we employed TurboID proximity labeling (Branon et al. 2018; Sanchez et al. 2021b). We generated a worm strain expressing V5::TurboID::COSA-1, enabling biotinylation of proteins within ∼10 nm of COSA-1 (Kim et al. 2014) (**Figure 2D**). Streptavidin signal was detected at CO-designated sites specifically in this strain, and mass spectrometry analysis identified COSA-2 among the proteins biotinylated by V5::TurboID::COSA-1, along with CDK-2 and the CO-associated protein RMH-1 (Jagut et al. 2016) (**Supplemental Figure S4**). Furthermore, affinity purification using streptavidin beads followed by immunoblotting showed specific enrichment of COSA-2::3xFLAG in a strain co-expressing both COSA-2::3xFLAG and V5::TurboID::COSA-1, but not in a strain expressing COSA-2::3xFLAG alone (**Figure 2D, Supplemental Figure 4C-E**).

Together, these data establish COSA-2 as a component of the meiotic CO machinery that becomes concentrated in late meiotic prophase at the sites of eventual CO formation and is in close physical proximity to the COSA-1–CDK-2 complex.

### COSA-2 forms foci specifically at CO-designated sites

While COSA-2 colocalizes with other CO factors at late-prophase CO-designated meiotic DSBR sites, the onset of its arrival is distinct. To facilitate visualization of recombination factors at early DSBR sites, we used a nuclear spread preparation that releases soluble protein pools, thereby improving detection of chromatin-bound proteins (**Figure 3**) (Pattabiraman et al. 2017; Woglar and Villeneuve 2018). In contrast to foci of MutSɣ component MSH-5 and the BLM DNA helicase (HIM-6 in *C. elegans*), which mark multiple DSB repair intermediates per chromosome pair in early pachytene nuclei and become restricted to a single CO-designated site per chromosome pair in late pachytene nuclei (Yokoo et al. 2012; Jagut et al. 2016; Woglar and Villeneuve 2018), COSA-2 foci appear abruptly at the onset of late pachytene and colocalize with BLM and MSH-5 exclusively at the 6 CO-designated sites per nucleus (**Figure 3A**). This abrupt appearance of COSA-2 also contrasts with COSA-1, which localizes as numerous faint foci per nucleus prior to accumulating as 6 bright foci per nucleus (Woglar and Villeneuve 2018; Haversat et al. 2022). The timing of appearance of COSA-2 foci does not reflect absence of the COSA-2 protein at earlier stages, as live imaging of COSA-2::GFP expressed from the endogenous *cosa-2* locus indicates that COSA-2 is present in the nucleoplasm of all germline nuclei (**Figure 3B**). This implies that the concentration of COSA-2 into a focus is not governed by COSA-2 availability but rather by regulated progression of the meiotic prophase program. Previous work has provided evidence that robust pro-CO factor accumulation and CO site designation in late pachytene is driven in part by a positive feedback loop involving CDK-2–COSA-1-mediated phosphorylation of intrinsically-disordered C-terminal tails of MSH-5 and ZHP-3–ZHP-4 (Haversat et al. 2022; Zhang et al. 2025). In germ lines co-immunostained for COSA-2 and phosphorylated MSH-5 (MSH-5^pT1009^), we found that the intensities of the first detectable COSA-2 foci were strongly correlated with the intensities of the colocalized MSH-5^pT1009^ foci (**Figure 3C, Supplemental Figure S5**). This finding indicates that the onset of COSA-2 accumulation at recombination sites occurs concurrently with accumulation of MSH-5^pT1009^.

**Figure 3.**
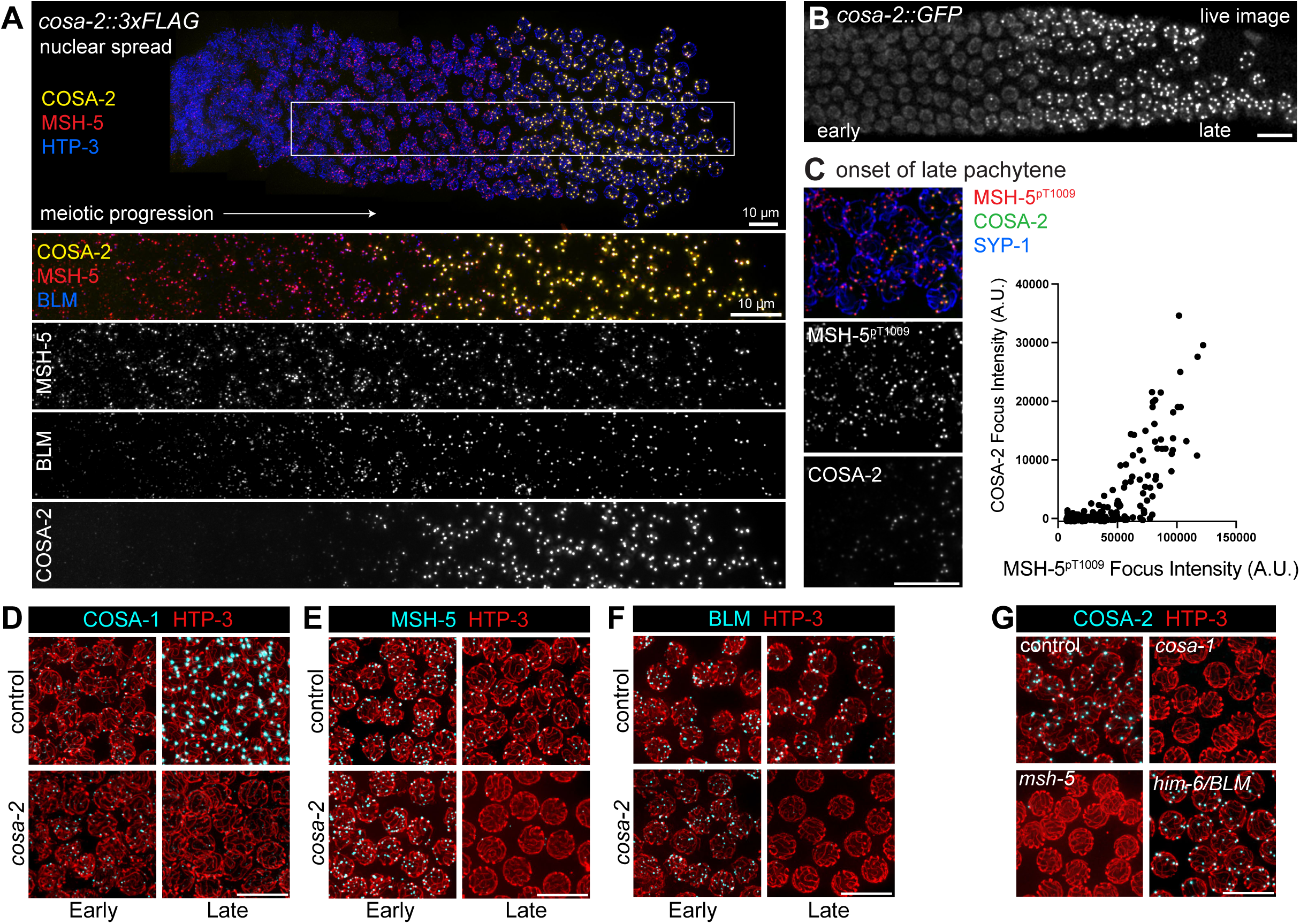
Colocalization and interdependencies of COSA-2 and other CO factors. (A) Nuclear spread preparation of *cosa-2::3xFLAG; him-6(blm)::HA* gonad that maintains spatiotemporal progression of meiotic prophase, illustrating localization dynamics of COSA-2::3xFLAG compared to MSH-5 and BLM::HA. HTP-3 immunostaining visualizes chromosome axes to aid in distinguishing individual nuclei. MSH-5 and BLM foci are detected throughout early and late pachytene stages, while COSA-2 foci appear abruptly at the onset of late pachytene, colocalizing with both MSH-5 and BLM. (B) Image of live GFP fluorescence in a dissected *cosa-2::gfp* germ line, revealing the presence of COSA-2::GFP in the nucleoplasm before it localizes as foci. (C) Left: nuclei from the region of the germ line where COSA-2 foci first appear, co-immunostained for phosphorylated MSH-5 (MSH-5^pT1009^) and COSA-2::FLAG. Right: plot depicting quantitative relationship between the peak intensities of individual MSH-5^pT1009^ foci (n = 212) and their colocalized COSA-2::3xFLAG signals in the represented field (A.U. = arbitrary units). (D-F) Immunolocalization of GFP::COSA-1, MSH-5, and BLM::HA in control and *cosa-2* mutant germlines, illustrating the retention of foci in early pachytene nuclei and loss of foci from late pachytene nuclei in the *cosa-2* mutant. (G) Immunolocalization of COSA-2::3xFLAG in control, *cosa-1*, *msh-5*, or *him-6/BLM* mutant backgrounds. All scale bars = 10μm.

### COSA-2 maintains CO factors at late pachytene recombination sites

Immunostaining for CO factors in *cosa*-2 mutant germ lines indicates that COSA-2 is required to maintain CO factors at late pachytene recombination sites. COSA-1 immunostaining revealed that numerous faint COSA-1 foci are visible in early pachytene nuclei in *cosa-2* mutants, but no bright COSA-1 foci are visible in late pachytene nuclei (**Figure 3D, Supplemental Figure S5**). Similarly, MSH-5 and BLM foci are still detected as multiple foci in early pachytene nuclei in *cosa-2* mutants, but MSH-5 foci and BLM foci are missing from late pachytene nuclei (**Figure 3E,F, Supplemental Figure S5**). The localization patterns of COSA-1, MSH-5, and BLM in *cosa-2* mutants indicate that COSA-2 is dispensable for loading of CO factors at early DSB repair intermediates, but is essential to maintain CO factors at late pachytene DSB repair sites. We refer to these COSA-2-scaffolded assemblages of CO factors at late pachytene DSB repair sites as “CO-specific repair compartments”.

Reciprocal immunolocalization experiments (**Figure 3G, Supplemental Figure S5**) showed that formation of COSA-2 foci is dependent on both COSA-1 and MSH-5, reflecting interdependence among these factors for recruitment to and/or maintenance at late pachytene recombination sites. However, as observed previously for COSA-1 foci, COSA-2 foci are not dependent on BLM, consistent with prior evidence that *BLM/him-6* mutant meiotic nuclei are competent for CO designation, but are defective for converting a subset of CO-designated repair intermediates into interhomolog COs (Wicky et al. 2004; Agostinho et al. 2013; Schvarzstein et al. 2014; Hong et al. 2016; Jagut et al. 2016).

### Evidence that COSA-2 functions after CO designation to ensure CO formation

In analyzing images of *cosa-2* mutant germ lines immunostained for MSH-5, we noted the presence of some nuclei with 6 MSH-5 foci in the region preceding the eventual complete loss of MSH-5 foci observed in later pachytene nuclei (**Figure 4A**). Systematic quantification of numbers of MSH-5 foci per nucleus during meiotic progression revealed a plateau of nuclei containing 6 MSH-5 foci before the total loss of MSH-5 foci occurring in *cosa-2* mutants (**Figure 4A**). Moreover, assessment of numbers of MSH-5 foci associated with individual resolvable HTP-3 tracks (in *cosa-2* mutant nuclei with 6 MSH-5 foci) revealed that nearly all synapsed chromosome pairs contained exactly one MSH-5 focus (97.9%, n = 144 HTP-3 tracks from 38 nuclei), as observed in nuclei from comparable regions of control germ lines (99.1%, n = 110 HTP-3 tracks from 38 nuclei) (**Figure 4B**). The presence of a temporarily sustained region of nuclei with a single MSH-5 focus per chromosome pair in *cosa-2* mutant germ lines was an initial indicator that COSA-2 is not required for the selection of a subset of early recombination intermediates as CO-designated sites.

**Figure 4.**
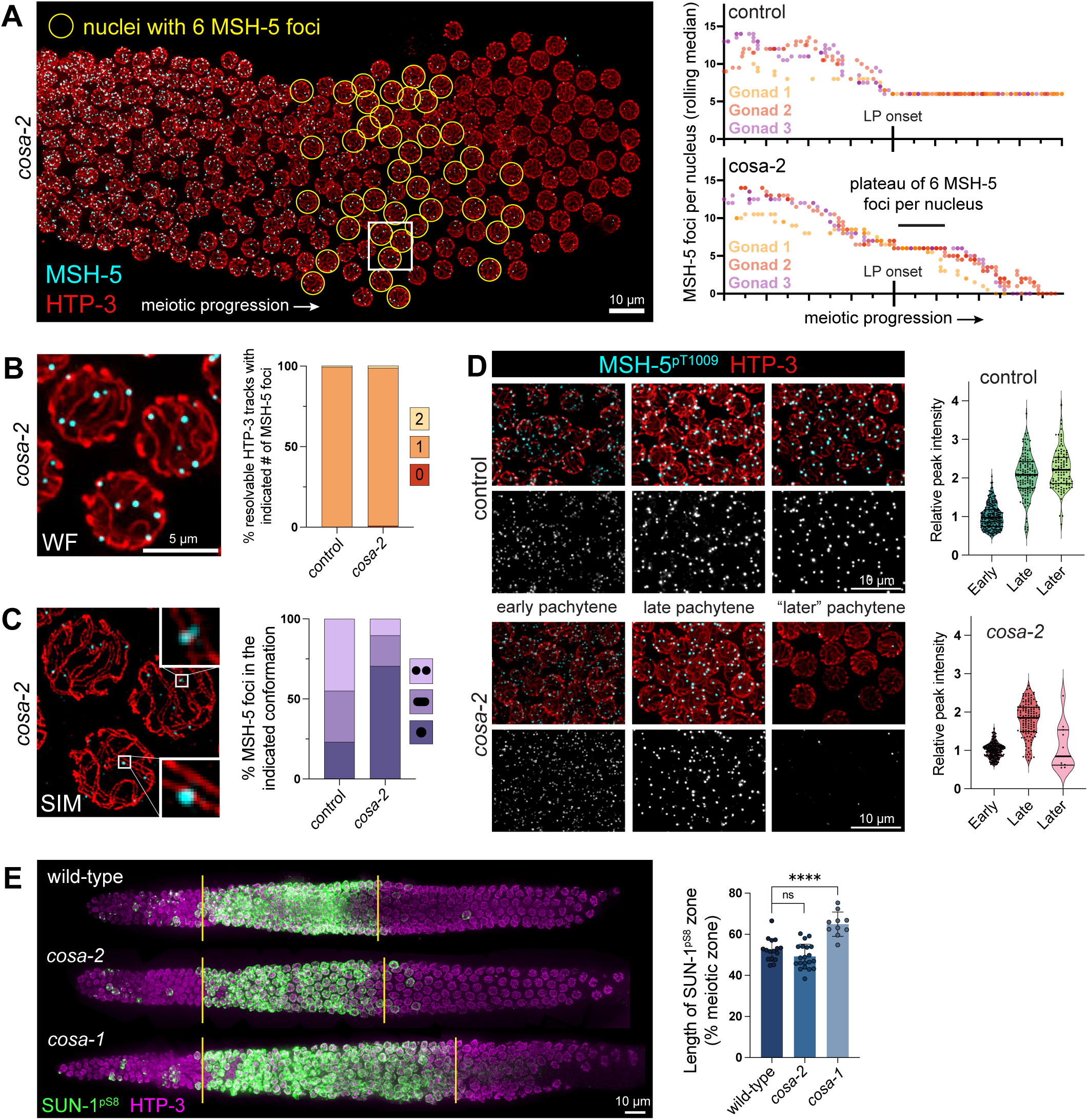
Evidence that COSA-2 functions after CO designation. (A) Left: Nuclear spread *cosa-2* mutant germ line illustrating decline in numbers of MSH-5 foci during progression from early pachytene to late pachytene. Nuclei containing 6 MSH-5 foci are circled in yellow. Right: Quantification of MSH-5 foci per nucleus through meiotic progression in nuclear spread germ lines, revealing a plateau of nuclei containing 6 MSH-5 foci before total loss of foci in the *cosa-2* mutant. Each data point reflects the median of 10 consecutive nuclei. The onset of late pachytene is defined as the first position in which the rolling median is 6 MSH-5 foci per nucleus for at least 3 consecutive measurements. (B) Left: inset corresponding to white box in A, representative wide field image of nuclei containing one MSH-5 (cyan) focus per chromosome pair. Right: quantification demonstrating that nearly all chromosome pairs contained a single MSH-5 focus in both control (109/110 chromosomes) and *cosa-2* mutant (141/144 chromosomes). (C) Left: Structured Illumination Microscopy (SIM) image of the nuclei in B, showing examples of MSH-5 foci (cyan) exhibiting the doublet (top inset) or singlet (bottom inset) conformations along parallel chromosome axes (HTP-3, red). Right: quantification of doublet, elongated, or singlet conformations (in nuclei containing 6 MSH-5 foci); for control, n = 147 foci scored, for *cosa-2,* n = 146 foci scored. (D) Left: representative fields of nuclei from spread germlines immunostained for MSH-5^pT1009^ and HTP-3. Right: quantification showing an increase in peak intensity of anti-MSH-5^pT1009^ staining from early pachytene to late pachytene in both control and *cosa-2* mutants. (E) Left: digitally-straightened images of whole mount of gonads immunostained for SUN-1^pS8^ as a proxy for CHK-2 activity and the status of the crossover assurance checkpoint. Right: quantification of the length of SUN-1^pS8^-positive (CHK-2 active) zone, showing no significant difference between wild-type (52.4%, n = 16) and *cosa-2* (49.2%, n = 20), but significant extension of the SUN-1^pS8^-positive zone in *cosa-1* (64.9%, n = 11). Statistical significance assessed via Mann-Whitney test: ns, P > 0.05; ****, P< 0.0001.

We used structured illumination microscopy (SIM) to examine the spatial architecture of the recombination sites in the transient “6 MSH-5 foci” nuclei from *cosa-2* mutant germ lines (**Figure 4C)**. In late pachytene nuclei from control worms, MSH-5 foci are predominantly detected as doublets or elongated foci (44.9% and 32.0%, respectively, n = 147 foci), conformations interpreted to reflect two populations of MutSɣ complexes embracing and stabilizing the DNA duplexes within an underlying double holiday junction (dHJ) CO-specific recombination intermediate (**Figure 4C**) (Woglar and Villeneuve 2018). Foci with the doublet and elongated conformations were also detected in “6 MSH-5 foci” nuclei from the *cosa-2* mutant, indicating that at least a subset of DNA break repair sites have progressed to the dHJ state and achieved a CO-specific repair intermediate conformation. However, the fraction of MSH-5 foci exhibiting the doublet or elongated conformation (19.2% and 10.3%, respectively; n = 146 foci) was reduced relative to control. The predominance of the MSH-5 singlet conformation in *cosa-2* mutants indicates that COSA-2 is required to fully establish and/or maintain CO-site architecture.

We also evaluated fluorescence signal intensity of phosphorylated MSH-5 as a proxy for phosphorylated MSH-5 accumulation, an alternative indicator of successful CO designation (Haversat et al. 2022) (**Figure 4D**). In control germ lines, peak immunofluorescence intensities of MSH-5^pT1009^ foci increase by roughly 2-fold between early pachytene and late pachytene and remained elevated through the end of late pachytene. Peak intensities of MSH-5^pT1009^ foci in the *cosa-2* mutant exhibited a similar 1.8-fold increase upon transition between early pachytene and the transient late-pachytene “6 MSH-5 foci” state, providing additional corroboration for the inference that *cosa-2* mutants are proficient for CO designation (**Figure 4D, Supplemental Figure S6**).

Further evidence that COSA-2 functions after CO designation is provided by experiments assessing the activity state of protein kinase CHK-2, a master regulator of meiotic progression that promotes multiple early prophase events including pairing, synapsis and DSB formation (MacQueen and Villeneuve 2001; reviewed in Kim et al. 2015) (**Figure 4E**). CHK-2 becomes activated at the onset of meiotic prophase, and its activity state can be visualized by immunostaining for nuclear envelope protein SUN-1 phosphorylated at Serine 8 (SUN-1^pS8^)(Penkner et al. 2009). Exit from the early pachytene stage is coupled to loss of SUN-1^pS8^ signal, which serves as a marker that all chromosome pairs within a nucleus have successfully formed CO-eligible recombination intermediates, thereby satisfying the “CO assurance checkpoint” (Rosu et al. 2013; Stamper et al. 2013; Woglar et al. 2013; Kim et al. 2015; Yu et al. 2016; Zhang et al. 2023). The CHK-2 active (SUN-1^pS8^-positive) zone is extended in germlines of mutants that are defective in forming CO intermediates, which consequently fail to satisfy the checkpoint, as illustrated here for the *cosa-1* mutant (Rosu et al. 2013; Woglar et al. 2013) **(Figure 4E**). In contrast, the CHK-2 active zone is not extended in *cosa-2* mutant germ lines (**Figure 4E**), indicating timely satisfaction of the CO assurance checkpoint. These data support the conclusion that *cosa-2* mutants are proficient for designation and formation of CO intermediates but are defective in converting these intermediates into COs.

### COSA-2 is required to constrain late-prophase localization of ZHP-3

As discussed above, *C. elegans* meiotic bivalents become asymmetrically reorganized around the presumptive CO site during meiotic prophase progression, with SC central region proteins and a subset of meiotic axis proteins becoming progressively enriched or depleted from short and long arm domains, beginning near the end of the pachytene stage (Nabeshima et al. 2005; Martinez-Perez et al. 2008). During this process of short-arm/long-arm differentiation, CO factor ZHP-3 likewise becomes progressively constrained in its localization: ZHP-3 is distributed throughout the length of the SCs in early pachytene, begins to become co-enriched with the SYP proteins on the short arms during late pachytene, and gradually becomes co-concentrated with other CO factors at the CO site as nuclei progress to the diplotene stage (Jantsch et al. 2004; Bhalla et al. 2008; Yokoo et al. 2012; Zhang et al. 2018). This progressive relocalization of SYP-1 and ZHP-3 is readily observed in control germ lines immunostained for HTP-3, SYP-1 and ZHP-3, resulting in a characteristic ZHP-3 “comet head” / SYP-1 “comet tail” localization pattern in diplotene nuclei (**Figure 5A**). Relocalization of SYP-1 and ZHP-3 is less prominent in *cosa-2* mutant germ lines, with both proteins remaining more broadly distributed within the SCs through the end of pachytene (**Figure 5A**). As chromosomes desynapse in the diplotene stage, SYP-1 does eventually become enriched on a sub-domain of the chromosome axes, likely an outcome of a symmetry breaking event triggered by CO designation, but SYP-1 depletion elsewhere appears less complete in *cosa-2* mutants. Moreover, ZHP-3 no longer becomes concentrated to a focal “comet head”, and instead colocalizes broadly with the enriched SYP-1 signal. (**Figure 5A**). These data indicate COSA-2 is required for ZHP-3 to become locally concentrated into a CO-site compartment.

**Figure 5.**
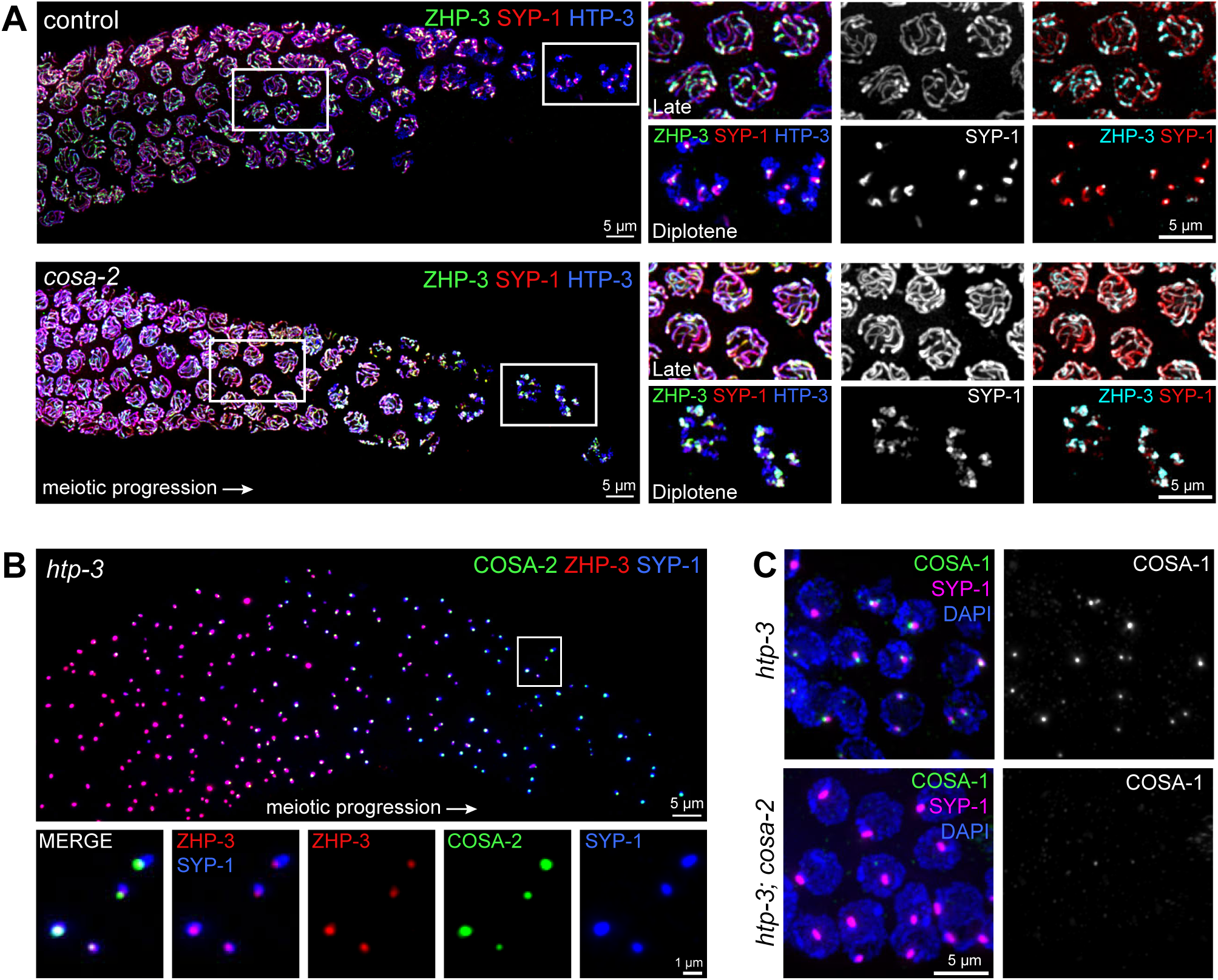
COSA-2 promotes appropriate ZHP-3 localization and organization of ectopic CO factor compartments. (A) Images of late pachytene through diplotene regions of whole mount germ lines co-immunostained for ZHP-3, SYP-1 and HTP-3, with insets highlighting ZHP-3 and SYP-1 localization in late pachytene and diplotene nuclei. The focal “comet head” enrichment of ZHP-3 that is normally observed at the diplotene stage was lost in the *cosa-2* mutant, and ZHP-3 was instead distributed throughout the entirety of the SYP-1 enriched domain. Other distinctions between control and *cosa-2* images are described in Results. (B) Image of polycomplexes in the late pachytene region of a whole mount germ line from an *htp-3; cosa-2::3xFLAG* worm, with insets showing COSA-2::3xFLAG co-localizing with ZHP-3 in a sub-domain of the polycomplex (marked by SYP-1). (C) Image of polycomplexes in late pachytene nuclei immunostained for COSA-1::GFP and SYP-1, showing that COSA-1 fails to localizes to polycomplexes in the *cosa-2* mutant.

We also examined COSA-2 localization and function in the context of ectopic CO-factor rich compartments (**Figure 5B,C**). Specifically, we examined germ lines from *htp-3* mutant worms, in which lack of axial elements results in SC-central region proteins forming nuclear aggregates called polycomplexes (1-2 per nucleus). This system enables testing of *in vivo* interactions and interdependencies among CO factors, as these factors have been observed to accumulate in sub-compartments of polycomplexes in the absence of underlying DSB repair intermediates (Goodyer et al. 2008; Severson et al. 2009). Previous work describes ZHP-3 as initially being distributed throughout the polycomplex (i.e. colocalizing with the majority of the SYP-1 signal), then later becoming restricted to a small focus at the periphery of the polycomplex that colocalizes with other CO factors (i.e. COSA-1 and MSH-5), paralleling ZHP-3 localization dynamics during wild-type meiotic prophase progression (Jantsch et al. 2004; Zhang et al. 2018; Köhler et al. 2025). COSA-2 likewise colocalizes with ZHP-3 foci at the periphery of polycomplexes during later prophase in the *htp-3* mutant, identifying COSA-2 as an additional component of these ectopic CO-factor-rich compartments (**Figure 5B**). Further, COSA-2 is required for localization and accumulation of COSA-1 in peripheral polycomplex foci (**Figure 5C**), implicating COSA-2 in the formation and/or maintenance of these ectopic CO-factor-rich compartments.

### COSA-2 is required continuously to maintain CO compartments

Our data led to the working model that COSA-2 functions after CO designation to concentrate CO-promoting factors, to stabilize CO-site architecture, and to maintain CO-specific repair compartments, thereby enabling CO-specific intermediates to mature as CO products. To investigate a potential requirement for COSA-2 in ongoing maintenance of CO compartments, we employed the Auxin Inducible Degron (AID) system (Zhang et al. 2015; Ashley et al. 2021) to assess the consequences of degrading COSA-2 in the late pachytene stage after CO-specific repair compartments have already been established. By using anti-FLAG immunostaining to detect COSA-2, we determined that exposing *AID::cosa-2::AID::3xFLAG* worms to auxin for 2 hours is sufficient to deplete AID::COSA-2::AID::3xFLAG foci from late pachytene germline nuclei (**Figure 6A, Supplemental Figure S7**).

**Figure 6.**
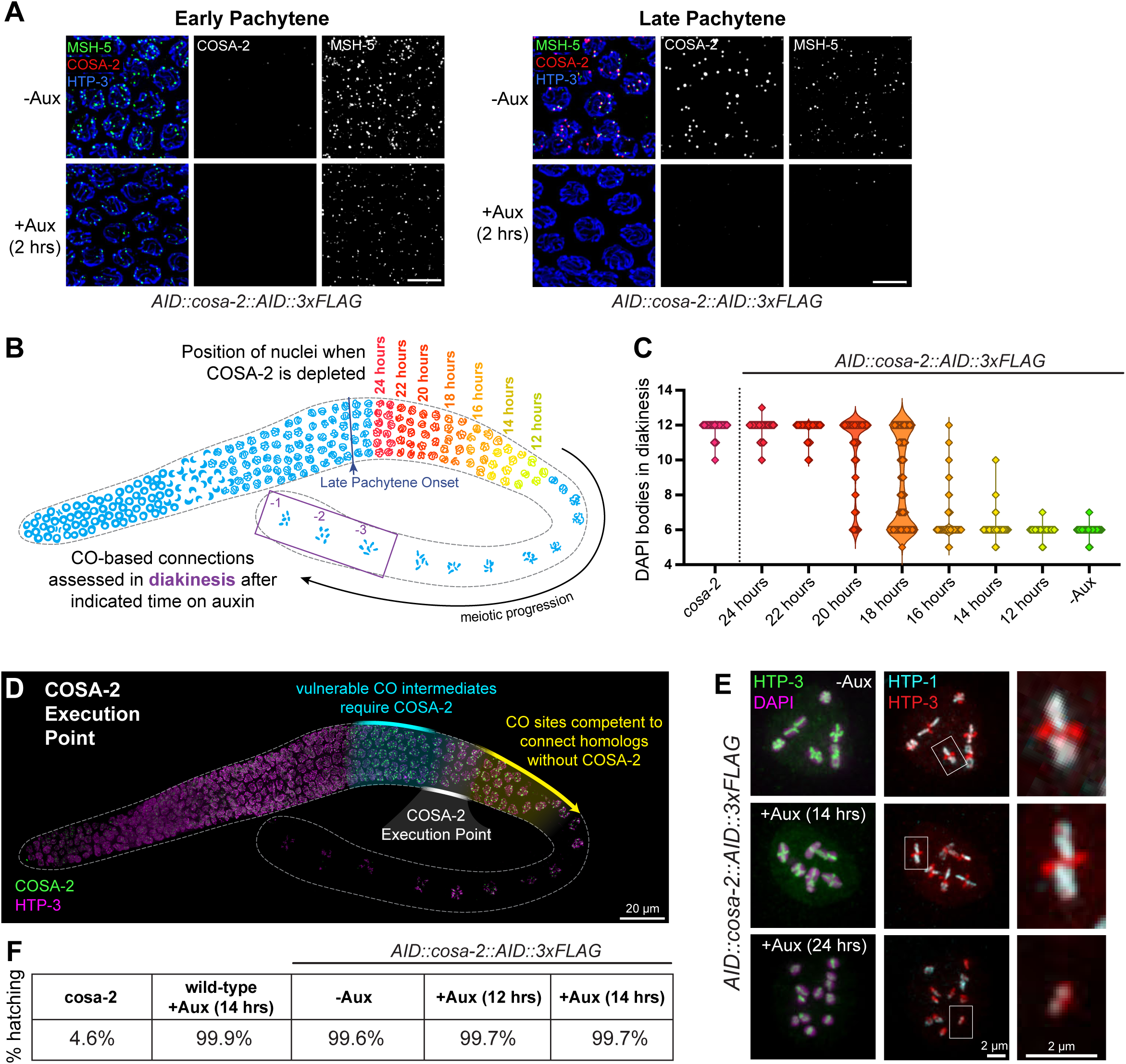
Identification and analysis of the COSA-2 execution point. (A) Images of early and late pachytene nuclei costained for AID::COSA-2::AID::3xFLAG and MSH-5, demonstrating robust depletion of COSA-2 foci from the late pachytene region after 2 hours of auxin exposure. Whereas early pachytene MSH-5 foci persist after auxin exposure, late pachytene MSH-5 foci are lost from late pachytene nuclei following auxin exposure, concomitant with loss of COSA-2 foci. (B) Schematic illustrating the experimental approach used to assess the competency of CO sites (in nuclei progressing through the late pachytene stage) to connect homologs in the absence of COSA-2. Success or failure to connect homologs was assessed in proximal diakinesis oocyte (purple box) following auxin exposures ranging from 12-24 h. For each indicated duration of auxin exposure, colors represent the approximate positions of scored nuclei at the time when COSA-2 was initially depleted. (C) Quantification of DAPI-stained bodies in proximal diakinesis nuclei after the indicated times on auxin. The 18-hour auxin exposure time point (n = 321 diakinesis nuclei) corresponds to nuclei that had been in the middle of late pachytene when COSA-2 was depleted (see Schematic) and marks a transition between nuclei containing mostly univalents (24-20 hours, n = 477, 116, 116 respectively) and nuclei containing mostly bivalents (16-12 hours, n = 293, 295, 230, respectively). (D) Schematic of inferred COSA-2 execution point overlayed on a whole-mount germline immunostained for HTP-3 and COSA-2::3xFLAG. Cyan represents a region containing nuclei in which CO-designated recombination intermediates that lose COSA-2 fail to mature as COs (as assessed by bivalent formation in diakinesis). Yellow indicates a region of nuclei that no longer require COSA-2 to establish or maintain crossover-based connections or to produce properly differentiated bivalents capable of undergoing reliable segregation. The COSA-2 execution point (in the middle of the late pachytene stage, indicated in white) represents the transition between these two states, *i.e.* the time during which COSA-2 completes its function and becomes dispensable for a successful outcome of meiosis. (E) Images of chromosomes from diakinesis oocytes immunostained for HTP-3 and for HTP-1, an axis component specifically enriched on the long arms of the bivalents, demonstrating that bivalents from nuclei in which COSA-2 had been depleted during the end of late pachytene (*i.e.* after the execution point) exhibit normal short arm/long arm differentiation. (F) Quantification of hatching rates for embryos laid during a 4-hour window following a 12 or 14 hour auxin treatment.

Notably, depletion of COSA-2 by a 2 hour auxin exposure resulted in a loss of integrity of CO-site compartments. MSH-5 foci were lost from late pachytene nuclei concomitantly with loss of COSA-2 foci, whereas early pachytene MSH-5 foci (which are not COSA-2 dependent) were retained (**Figure 6A**). A 2 hour auxin treatment likewise resulted in loss of focal enrichment of ZHP-3 in “comet heads” at the diplotene stage (**Supplemental Figure S7**), and ZHP-3 was instead distributed throughout the entirety of the SYP-1 enriched domain.

The rapid disruption of CO factor localization upon COSA-2 depletion indicates that COSA-2 is required for ongoing maintenance of CO-site compartments and reinforces the idea that COSA-2 is critical for stabilizing CO factors at these sites.

### COSA-2 Execution Point Analysis

The ability to disrupt CO repair compartments after their initial assembly by conditional degradation of COSA-2 created a unique opportunity to conduct a COSA-2 execution point analysis, *i.e.* to determine the time during meiotic prophase when COSA-2 performs its essential function(s) (Hartwell et al. 1973). For this purpose, we exposed animals to auxin for durations ranging from 12-24 hours to assess diakinesis nuclei that had lost COSA-2 at different stages of progression through the late pachytene region (**Figure 6B, Supplemental figure S8**). In COSA-2-depleted nuclei that had progressed to the late diakinesis stage, presence or absence of CO-based connections (chiasmata) was assessed by counting DAPI-stained bodies (**Figure 6C**).

Multiple prior studies using a variety of approaches have established that nuclei progress through the pachytene stage at a rate of approximately one cell row per hour (Crittenden et al. 2006; Jaramillo-Lambert et al. 2007; Rosu et al. 2011; Hicks et al. 2022; Čavka et al. 2025) and that oocytes mature at a rate of approximately 1-2 per hour in young adult hermaphrodites of the age used in our analysis (McCarter et al. 1999). Based on this information, we estimate that nuclei take approximately 24 hours to progress from the onset of the late pachytene stage to late diakinesis. Thus, when scoring diakineses nuclei, we combined this progression timing estimate with the 2 hour auxin exposure required to deplete AID::COSA-2::AID::3xFLAG to infer the approximate position of the nuclei when COSA-2 was initially depleted (**Figure 6B,C**, Supplemental Figure S8).

Our analysis of the success of chiasma formation following COSA-2 depletion indicates that COSA-2 is required during the first half of the late pachytene stage to form CO-based connections between homologs, and enables several additional inferences. *AID::cosa-2::AID::3xFLAG* animals exposed to auxin for 24 hours had an average of 11.9 DAPI-stained bodies (n = 477 diakinesis nuclei), indicating that loss of COSA-2 at or soon after the onset of late pachytene results in failure to form chiasmata. 22 and 20 hours of auxin exposure resulted in an average of 11.8 (n = 116) and 10.6 (n = 116) DAPI-stained bodies respectively, indicating that COSA-2 continues to be required for the formation of CO-based connections for several more hours. 18 hours of auxin exposure resulted in an average of 8.8 DAPI-stained bodies (n=321), reflecting an increase in the number of chromosome pairs successfully connected via chiasmata. In animals exposed to auxin for shorter time periods, this shift to successful chiasma formation was complete: after 16, 14, and 12 hours on auxin, animals displayed an average of 6.2 (n=293), 6.0 (n = 295) and 6.0 (n = 230) DAPI-stained bodies, respectively. Thus, nuclei that have spent a longer time in the late pachytene stage before COSA-2 is depleted are competent for chiasma formation despite disruption of visible CO site compartments in these nuclei. These data identify a temporal-spatial window in the middle of the late pachytene stage wherein CO-specific repair intermediates transition from a vulnerable state requiring COSA-2 for completion of CO formation to a state where the local concentration of COSA-2 and associated CO factors at CO sites is no longer needed to connect homologs. We refer to this window as the “COSA-2 execution point” (**Figure 6D**). A COSA-2 execution point in the middle of the late pachytene stage indicates that the presence of COSA-2 during the first portion of late pachytene is sufficient to ensure CO formation.

While these data indicate that requirement for COSA-2 to connect homologs is completed before the end of the late pachytene stage, COSA-2 remains concentrated at CO/chiasma sites for much longer, persisting throughout the proximal (*i.e.* later) region of the late pachytene stage and the diplotene stage until the −3 or −4 diakinesis oocyte (**Figure 2A, 6D**). Thus, we investigated whether COSA-2 might have additional roles in promoting successful meiosis after the execution point defined above. First, we assessed whether depletion of COSA-2 after the execution point, *i.e.* from proximal late pachytene onward (14 h auxin exposure), would affect proper structural differentiation of late prophase meiotic bivalents. Specifically, we evaluated short-arm/long arm differentiation of diakinesis bivalents by immunostaining for HTP-3 and for HTP-1, an axis component that becomes enriched on the bivalent long arms (Martinez-Perez et al. 2008) (**Figure 6E**). Long-arm enrichment of HTP-1 on diakinesis bivalents following 14 h auxin exposure was indistinguishable from the no-auxin control, indicating that COSA-2 is not required during the proximal late pachytene, diplotene or diakinesis stages to achieve successful bivalent differentiation. We also assessed the viability of embryos laid during a 4-hour window following a 12 or 14 hour auxin treatment, as chromosome segregation errors during the ensuing meiotic divisions would result in unhatched inviable aneuploid embryos (**Figure 6F**). >99% of embryos from auxin-treated worms hatched and gave rise to viable progeny, indicating that the normal-appearing diakinesis bivalents present in oocytes in which COSA-2 had been depleted in late prophase are indeed functionally normal in supporting meiotic chromosome segregation.

Together, our data indicate that COSA-2 is required during the first half of the late pachytene stage to promote and ensure the maturation of CO-designated recombination intermediates into completed crossovers. After executing its essential function during this period, COSA-2 and the other CO factors maintained at CO sites by COSA-2 are no longer needed to connect homologs or to produce properly differentiated bivalents capable of supporting accurate chromosome segregation.

## DISCUSSION

This work establishes COSA-2 as a core component of the meiotic crossover machinery that is essential for CO maturation. COSA-2 specifically marks CO-designated sites and collaborates with other core CO-promoting factors including the COSA-1–CDK-2 complex, MutSɣ, BLM, and ZHP-3 to promote reliable CO formation. At a practical level, endogenously-tagged COSA-2 can serve as a robust marker for late pachytene CO-designated sites to aid future studies of CO compartments and CO patterning. At a conceptual level, this work provides new insights into the timing of CO formation, the composition and function of privileged CO-specific repair compartments, and the mechanisms ensuring reliable CO maturation.

### CO designation and CO maturation in *C. elegans* are genetically and temporally separable events

Defining features of COSA-2 are that: 1) it is not required until after CO designation, and 2) its localization to chromosomes occurs specifically at sites that have already been selected to become COs and at sites where CO maturation is actively occurring or has already been completed. These features distinguish COSA-2 from other known *C. elegans* CO-site associated factors, which either: a) appear as foci prior to the onset of late pachytene in numbers in that exceed the eventual number of COs, or b) are broadly distributed throughout the SC before eventually becoming enriched at CO sites.

The inference that COSA-2 is dispensable for CO designation is based on several lines of evidence. Numbers of MSH-5 foci in *cosa-2* mutants transiently plateau at 6 per nucleus (1 per chromosome pair) before disappearing. In these “6 MSH-5 nuclei”, a subset of foci exhibit a doublet conformation (implying the presence of CO-specific DNA intermediates (Woglar and Villeneuve 2018)) and/or an increase in MSH-5 phosphorylation (reflecting initiation of a feedback loop involved in robust CO designation (Haversat et al. 2022)). Moreover, *cosa-2* mutants exhibit timely satisfaction of the CO assurance checkpoint, indicating molecular recognition that all six chromosome pairs have acquired an intermediate that is competent to become a CO, and they also initiate symmetry-breaking events (i.e. bivalent short-arm/long-arm differentiation) normally associated with selection of a single CO site per chromosome pair. Collectively, these data indicate that selection of CO sites is successful in *cosa-2* mutants. Despite successfully achieving CO designation, *cosa-2* mutants nevertheless fail to mature CO-specific intermediates into CO products, emphasizing a clear distinction between CO designation and CO maturation. This mechanistic distinction between CO designation and CO maturation processes in *C. elegans* reinforces a conclusion of a recent report showing that mutants expressing modified COSA-1 proteins impaired for interactions with other CO factors exhibit defects in CO maturation despite apparent proficiency for CO designation (Yang et al. 2024).

In addition to being genetically separable, CO designation and CO maturation are temporally separable. Our execution point analysis reveals the earliest possible timing of crossover maturation to be in the middle of late pachytene (see below), at least 8 hours after COSA-2 foci are initially detected at CO-designated DSB repair sites. Further, live imaging of the dynamic behavior of COSA-1 foci by Čavka et al. (2025, BioRxiv) provides evidence that CO designation occurs ∼6 hours prior to entry into the late pachytene stage, long before COSA-2 “joins the party”.

Intriguingly, failure of CO-designated intermediates to reliably mature as COs has been proposed to underlie a substantial fraction of segregation errors in human meiosis. Wang et al. (2017) estimated that approximately one-quarter of CO-designated intermediates in human oocytes fail to mature as CO products, a phenomenon they refer to as “crossover maturation inefficiency”. They propose that crossover maturation inefficiency is a major contributor to oocyte-derived aneuploidy, compounding the effects of age-dependent loss of cohesin (reviewed in Wang et al. 2019; Gruhn and Hoffmann 2022). Crossover maturation inefficiency is likewise observed in spermatocytes of juvenile human and mouse males, but not adult males (Wang et al. 2017; Zelazowski et al. 2017), implying the presence of factors that modulate the efficiency of crossover maturation. In *C. elegans*, the central role of COSA-2 in ensuring reliable CO maturation provides an attractive opening to further explore the mechanisms governing and promoting reliable CO maturation.

### Function of COSA-2 in ensuring reliable CO maturation

Our data clearly demonstrate a requirement for COSA-2 in converting CO-designated recombination intermediates into functional connections that enable and ensure homolog segregation at the meiosis I division. Moreover, COSA-2 is required for maintaining the assemblages of CO-promoting factors at the sites where CO maturation takes place and for enriching the accumulation of factors that had initially been distributed through the SCs to those sites. How are these essential functions of COSA-2 accomplished?

Our thinking about how COSA-2 functions to promote CO maturation is informed by the abruptness of its recruitment to CO sites at late pachytene onset. Progression to late pachytene represents a developmental transition between an earlier cellular state during which nuclei accumulate DSB repair intermediates to a later state in which completion of repair is executed. This transition is triggered by satisfaction of the CO assurance checkpoint and consequent inactivation of the regulatory protein kinase CHK-2. A major challenge upon transition to the “completion of repair” stage is that repair must be completed differently at CO-designated sites versus non-CO sites present within the same nucleus. Woglar and Villeneuve (2018) proposed that this could be accomplished through creation of spatially-segregated compartments, which would enable distinct biochemistry to occur at CO and non-CO sites. We propose that the recruitment of COSA-2 at late pachytene onset promotes and reinforces this spatial compartmentalization.

When COSA-2 is not present to scaffold CO compartments, why might CO-designated intermediates fail to mature as CO products? At a molecular level, the absence of COSA-2 and the consequent breakdown of CO-factor-rich assemblages leads to the loss of dHJ stabilizing factors (*e.g.* MutSγ) at CO-designated sites and potentially exposes intermediates to factors such as dissolvase complexes or helicases with the ability to disassemble unprotected joint molecule intermediates to yield non-CO repair products (Youds et al. 2010; Jagut et al. 2016; Yang et al. 2024). Further, as COSA-1 was recently demonstrated to interact directly with the nuclease scaffolding protein HIM-18/SLX-4 to enable recruitment of structure-selective nucleases implicated in HJ resolution (Saito et al. 2009; Agostinho et al. 2013; O’Neil et al. 2013; Saito et al. 2013; Liu et al. 2026), the loss of COSA-1–CDK-2 complexes from these sites could also diminish local concentrations of HJ resolvase activities needed to execute CO-biased resolution. We propose that COSA-2 acts as a “molecular glue” to locally maintain large assemblages of factors that stabilize CO intermediates, positively regulate CO-promoting activities, and/or recruit resolvases. We further speculate that COSA-2 may also serve as “insulation” to prevent enzymes with non-CO promoting activities from gaining access to and dismantling CO intermediates within CO-specific repair compartments.

Our ability to conditionally degrade COSA-2 at different stages of late prophase progression and consequently dismantle existing CO compartments has defined a COSA-2 “execution point” after which concentration of CO factors at CO-designated sites is no longer required to connect homologs. It is tempting to speculate that this execution point reflects the timing when CO-specific intermediates (i.e. dHJs) mature into CO products via dHJ resolution, but we do not have a means to assess this directly. Nevertheless, we can infer that at the COSA-2 execution point in the middle of the late pachytene stage, the intermediates present at CO-designated DSB repair sites transition from a state where they are vulnerable to being dismantled (and repaired to yield non-CO products) to a non-vulnerable state that reliably results in CO-biased resolution. The vulnerable state (where DSB repair intermediates require COSA-2 and other CO factors for protection) lasts for approximately 8 hours after COSA-2 foci are initially detected at CO-designated sites at the onset of late pachytene. The ability to disassemble pre-existing CO-site compartments via COSA-2 degradation further implies that these compartments require active ongoing maintenance for their sustained integrity and function.

### Achieving Order through Disorder

Our discovery of COSA-2 as a largely unstructured protein responsible for maintaining *C. elegans* meiotic CO-specific repair compartments epitomizes a recurring theme of intrinsically-disordered protein domains contributing to the organization and function of the multi-protein mega-complexes that assemble at nascent CO sites. There are multiple examples in *C. elegans* alone. Appended to its highly-conserved MutS “body”, *C. elegans* MSH-5 has an intrinsically-disordered C-terminal tail that comprises ∼40% of the protein and plays critical scaffolding roles in recruitment of pro-CO factors, including the COSA-1–CDK-2 complex (Haversat et al. 2022). COSA-1 itself has a short N-terminal intrinsically disordered region (IDR) containing key residues required for interactions with both MSH-5 and ZHP-3 and for normal accumulation of COSA-1 at late pachytene CO-designated sites (Yang et al. 2024). Meiotic RING finger protein ZHP-3 has an extended C-terminal IDR that is sufficient to recruit the C-terminal IDRs of MSH-5 and related meiotic RING finger protein ZHP-4 to droplets when co-expressed in HEK293 cells (Zhang et al. 2025), and deletion of most of the ZHP-4 IDR compromises COSA-1 and ZHP-3 localization and CO formation *in vivo* (Nguyen et al. 2018). Related meiotic RING proteins have been identified as important CO-promoting factors in multiple organisms and exhibit dynamic association with SCs and/or recombination sites, *e.g.* Zip3 in budding yeast, HEI10 in *Sordaria*, *Arabidopsis* and mice, and RNF212 and RNF212B in mice (Agarwal and Roeder 2000; Ward et al. 2007; Chelysheva et al. 2012; Reynolds et al. 2013; De Muyt et al. 2014; Qiao et al. 2014; Condezo et al. 2024; Ito et al. 2025); these proteins similarly contain long IDRs comprising a substantial fraction of total protein length. Further, the HEI10 interacting protein HEIP1, which has been implicated in promoting/regulating CO formation in rice, Arabidopsis, and mice, harbors only a short ∼70 amino acid conserved domain and is predicted to be disordered over most of its length (Li et al. 2018; Singh et al. 2023; De Muyt et al. 2025). We speculate that for COSA-2 and other proteins involved in assembly, maintenance and/or function of CO-site compartments, the presence of large disordered stretches may provide conformational flexibility between protein interaction domains that enables simultaneous binding with multiple protein partners (Dunker et al. 2005).

Do orthologs or analogs of COSA-2 function in mammalian meiosis? Although BLAST analyses do not confidently identify COSA-2 orthologs outside the *Caenorhabditis* genus, our structural modeling using Alphafold reveals striking parallels between COSA-2 and mammalian PRR19, a disordered protein that is required for CO formation in mice and interacts with CNTD1, the mammalian COSA-1 ortholog (Bondarieva et al. 2020) (**Supplemental Figure S3**). Despite extremely limited sequence similarity, PRR19 is predicted to interact with CNTD1 using two alpha-helices in a manner remarkably similar to the predicted COSA-2–COSA-1 interaction interface. Superposition of the predicted cyclin-binding helices of COSA-2 and PRR19 bound to their respective cyclin partners shows that both proteins are predicted to engage nearly identical hydrophobic surfaces of COSA-1 and CNTD1, and key hydrophobic residues within the cyclin-binding helices are preserved across both protein families and are predicted to contact conserved hydrophobic residues on the cyclins. Notably, COSA-2 and PRR19 are both predicted to bury two full alpha-helices within an extended groove formed by the elongation of helix α2 and the insertion of helix α2.5, features unique to the COSA-1/CNTD1 cyclin subfamily (Yokoo et al. 2012). The conservation of these structural elements in COSA-1 and CNTD1 and the corresponding groove-binding architecture of COSA-2 and PRR19 supports the co-evolution of these protein pairs. Thus, our findings regarding the functional roles of COSA-2 are likely relevant to mammalian meiosis and can elevate our understanding of how we make gametes and embryos with the appropriate number of chromosomes.

## MATERIALS AND METHODS

### *C. elegans* maintenance, strains, and gene editing

Strains used in this study were cultured on nematode growth medium (NGM) agar plates at 20 °C according to standard conditions (Brenner 1974; Stiernagle 2006) unless otherwise stated. For strain details, see **Tables 1 and 2**. For experiments involving strains containing balancer chromosomes, homozygous mutant worms were identified by lack of markers associated with the balancer chromosome and synchronized by selection as L4 larvae.

**Table 1:**
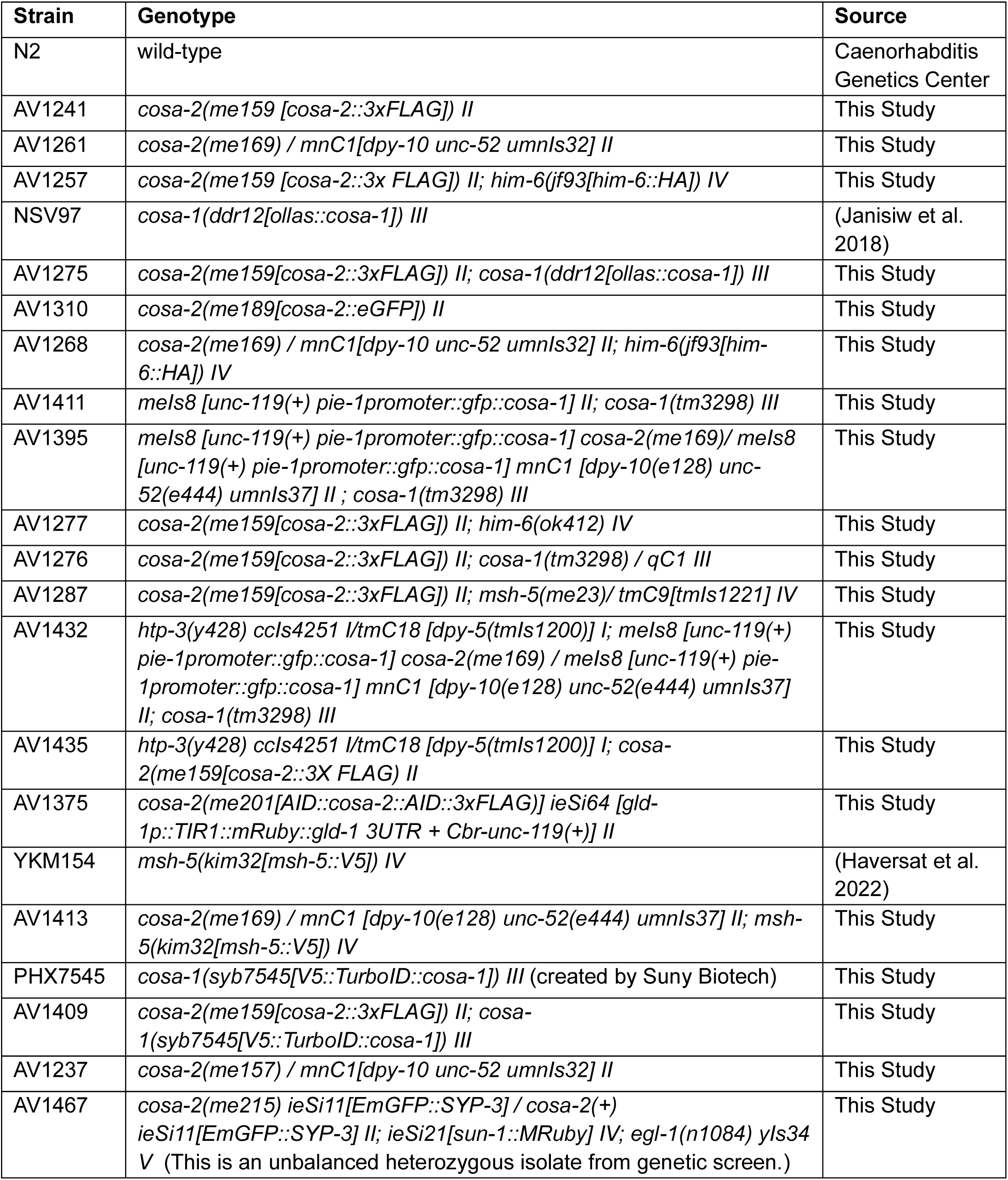
Strains used in this study. The following *C. elegans* strains were used in this study and are available from AMV upon request.

**Table 2:** Alleles generated in this study.

| **Allele** | **Genotype** | **Mutagenesis Information** |
| --- | --- | --- |
| *me157* | *cosa-2(me157) II* | CRISPR modification of N2 wild-type to introduce II: 8,275,739 C -> T (splice acceptor mutation) |
| *me159* | *cosa-2(me159[cosa-2::3xFLAG]) II* | CRISPR modification of N2 wild-type |
| *me169* | *cosa-2(me169) II* | CRISPR modification of N2 wild-type to introduce STOP-IN (Wang et al. 2018) into Exon 2 |
| *me188* | *cosa-2(me188[cosa-2::eGFPfragment]) II* | CRISPR modification of N2 wild-type to introduce nested GFP fragment (Vicencio et al. 2019) |
| *me189* | *cosa-2(me189[cosa-2::eGFP]) II* | CRISPR modification of *cosa-2(me188[cosa-2::eGFPfragment]) II* |
| *me194* | *cosa-2(me194[cosa-2::AID::3xFLAG]) II* | CRISPR modification of *cosa-2(me159[cosa-2::3xFLAG]) II* |
| *me201* | *cosa-2(me201[AID::cosa-2::AID::3xFLAG]) II* | CRISPR modification of *cosa-2(me194[cosa-2::AID::3xFLAG]) II* |
| *syb7545* | *cosa-1(syb7545[V5::TurboID::cosa-1]) III* | CRISPR modification of N2 wild-type |
| *me215* | *cosa-2(me215) II* | EMS-induced mutation<br>II: 8,274,596 A -> T (Leu428* early stop) |

New alleles of *cosa-2* (previously *gei-14*) were generated by CRISPR-based genome engineering as previously described (Arribere et al. 2014; Paix et al. 2015). Guide RNAs and PAM sites for all targets were designed with the IDT “Custom Alt-R CRISPR- Cas9 Design Tool”. In brief, an injection mixture was assembled by adding tracrRNA (IDT), *dpy-10* crRNA (co-injection marker, (GCTACCATAGGCACCACGAG, IDT), and target crRNA. The solution was boiled at 95 °C for 5 minutes and allowed to cool to room temperature. Cas9 (CP02, PNA Bio), *dpy-10* co-injection marker repair ssODN, target repair oligo (IDT Ultramer or PCR amplicon, see **Table 3**), and Duplex Buffer (to a total volume of 10 µL) were added and the solution was spun at max speed and loaded into a micropipette for injection into the germ line. Homozygous edits were identified by PCR screening using the primers in **Table 3** and confirmed by Sanger sequencing. For edits occurring on the same chromosome as the *dpy-10* co-injection marker (i.e. *cosa-2* on chromosome *II*), morphologically wild-type progeny of injected worms were isolated from “jackpot” plates (indicated by an abundance of Rol and Dpy animals) and subjected to PCR screening and Sanger sequencing to detect edits.

**Table 3:**
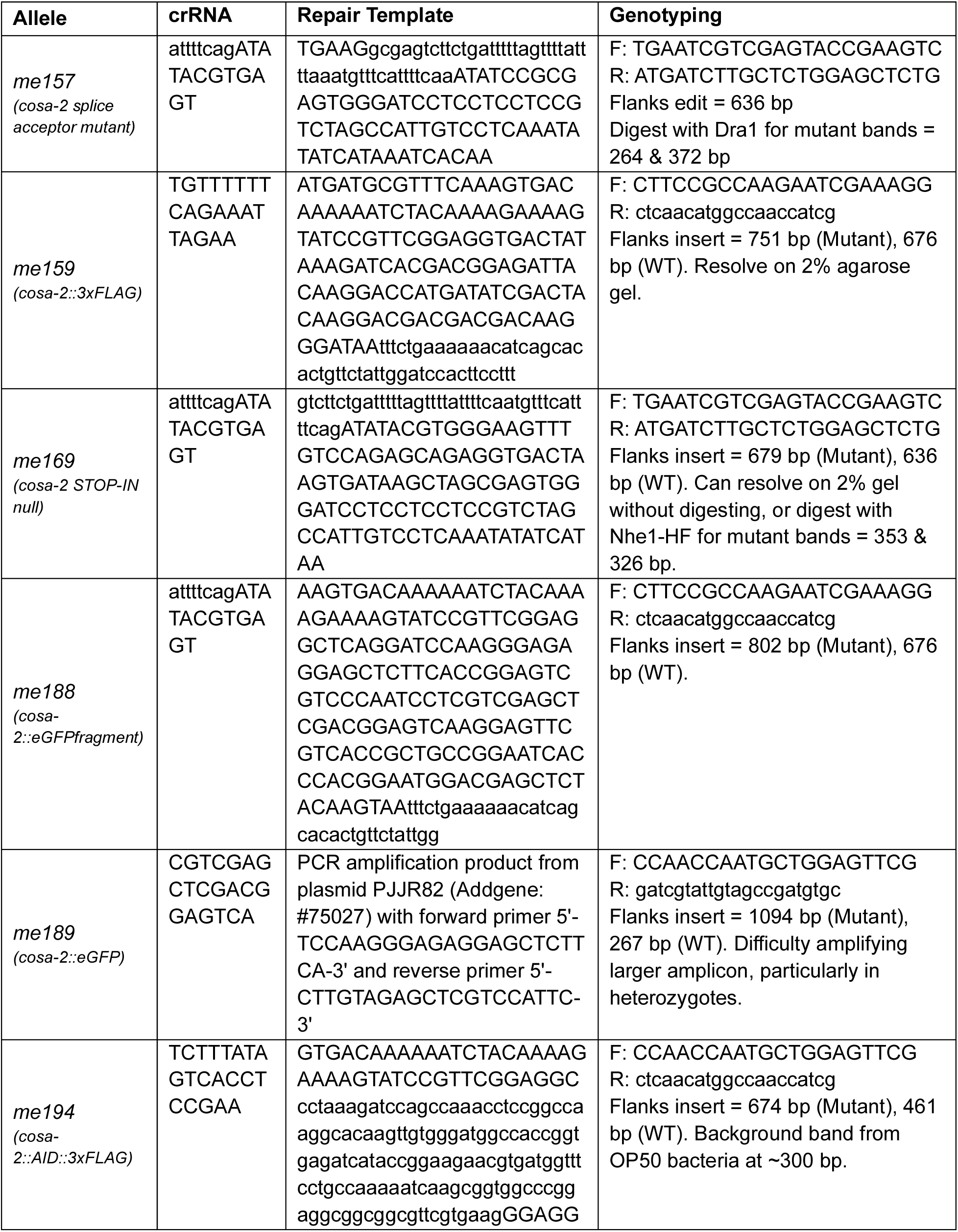
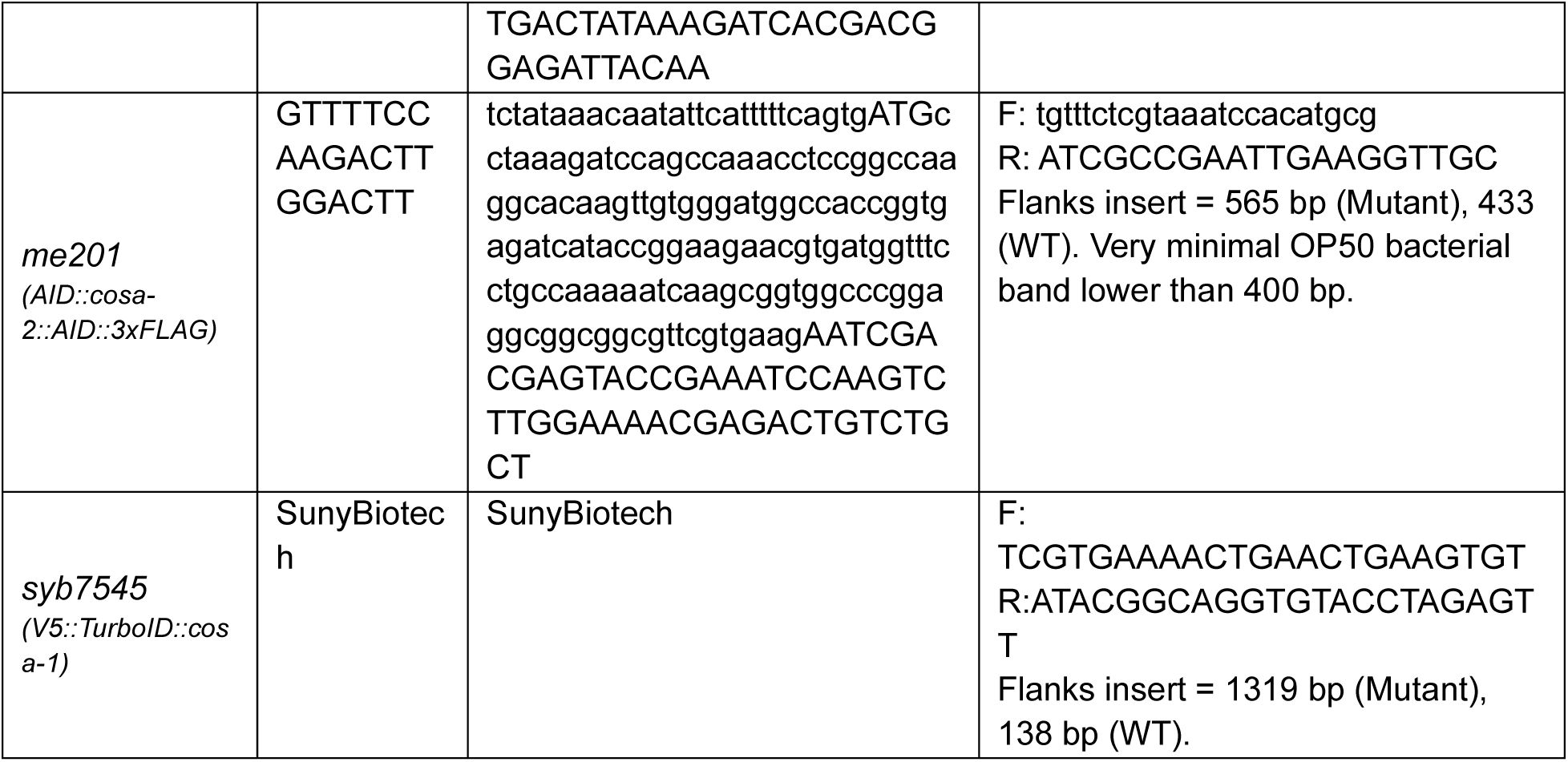
crRNAs, repair templates, and genotyping primers for mutant alleles generated in this study.

### Analysis of progeny viability and incidence of males

Single L4 hermaphrodites (5 animals per genotype in 2 independent replicates) were placed on individual plates. Each parent was transferred to a new plate at 24 hour intervals until 72 hours post L4. Number of unhatched eggs were counted 24 hours after plating the parent worm on each plate. Number of hatched progeny were counted 72 hours after plating the parent worm on each plate. “Total Eggs Laid” is the sum of unhatched eggs plus the total number of live counted progeny. Percent hatching = (live progeny/total eggs) x 100. Percent males = (total males/live progeny) x 100. Viability assessment for Figure 7 was similarly quantified, but with 12 replicate plates in which cohorts of 4 worms were permitted to lay eggs for 4 hours.

### DAPI staining and quantification

4’, 6-Diamidino-2-phenylindole (DAPI) staining of whole animals was performed on adult worms 24-26 hours post-L4 larval stage as a means to quantify success or failure of bivalent formation in oocyte nuclei, using a procedure modified from Bessler et al. (2007). Animals were picked into a drop of M9 buffer on a glass slide and agitated to wash away bacteria. Excess buffer was aspirated and replaced with 30 µL of 100% Ethanol. Samples were air-dried until worm carcasses were fully desiccated. Ethanol treatment was repeated in cases of insufficient desiccation. 7 µL of 5 µg/mL DAPI in M9 buffer was added to rehydrate the worms. 7 µL of 5 µg/mL DAPI in Vectashield was added as mounting medium. Samples were mounted, sealed with nail polish, and visualized within 2 weeks to avoid DAPI fading. DAPI-stained body counts were performed on a Zeiss Axio ImagerM2 by manually counting resolvable DAPI bodies in the 3 most proximal oocyte nuclei while through-focus scanning with a 63x oil objective. Both the anterior and posterior germline of 10-20 animals were assessed for each replicate, treatment, and/or genotype.

### Cytological Preparation and Immunostaining

**Whole mount germline preparation and immunostaining** was performed as in Phillips et al. (2009), with modifications. Briefly, adult animals 24-26 hours-post-L4 were scalpel dissected in egg buffer containing 0.1% Tween-20 and fixed with 1% v/v EM-grade formaldehyde solution for 5 minutes. Animals were permeabilized by freeze-cracking and further fixed in ice-cold 100% methanol for 2 minutes. Slides were washed 3 x 10 minutes in PBST (1x PBS, 0.1% Tween) followed by blocking in 1% BSA in PBST with 0.05% w/v sodium azide for 1 hour. Wash steps were repeated before and after primary and secondary antibody application. Slides were counterstained with DAPI by adding 5 µg/mL DAPI in the penultimate wash. Slides were mounted with Vectashield and sealed with nail polish.

**Nuclear spread preparations** were performed as described in Pattabiraman et al. (2017) and Woglar and Villeneuve (2018). At least 100 worms were rapidly dissected in a small volume (<10 µL) of 10% v/v Hanks Balanced Salt solution (StemCell Technologies) plus 0.1% Tween-20 in water. 50 µL of spreading solution was distributed over the coverslip region of the slide [Spreading solution per slide: 32 μl of Fixative (4% w/v Paraformaldehyde and 3.2–3.6% w/v Sucrose in water), 16 μl of Lipsol solution (1% v/v/ Lipsol (obtained from Josef Loidl) in water), 2 μl of Sarcosyl solution (1% w/v of Sarcosyl in water)]. Slides were dried undisturbed overnight at room temperature and then immersed in ice cold 100% methanol for 20 minutes. Immunostaining and slide mounting was then performed as described above.

**Antibodies** in this study include the following primary antibodies used at the indicated dilutions for immunofluorescence: Chicken anti-HTP-3 (1:200 or 1:500) (MacQueen et al. 2005 and see below), rabbit anti-GFP (1:1,500) (Yokoo et al. 2012), guinea pig anti-ZHP-3 (1:500) (Bhalla et al. 2008), guinea pig anti-SYP-1 (1:200) (MacQueen et al. 2002), rabbit anti-SYP-1 (1:250) (MacQueen et al. 2002), rabbit anti-RAD-51 (1:500) (Colaiácovo et al. 2003), affinity purified rabbit anti-MSH-5pT1009 (1:25) (Haversat et al. 2022), rat anti-HIM-8 (1:500) (Phillips et al. 2005), rabbit anti-HTP-1/2 (1:500) (Martinez-Perez et al. 2008), rabbit anti-MSH-5 (1:10,000) (SDI/Novus 3875.00.02), rat anti-HA (1:2,000) (Roche 11 867 423 001), mouse anti-FLAG (1:2,000) (Sigma-Aldrich F1804), rabbit anti-OLLAS (1:1,500) (Genscript A01658), Streptavidin Alexa Fluor 647 conjugate (1:1000) (Invitrogen), and GFP-booster Alexa Fluor 488 (1:200) (Proteintech). Secondary antibodies conjugated to Alexa dyes 405, 488, 555, or 647 were used at 1:200, 1:500, or 1:1000 dilutions. The following antibodies were used for immunoblotting (Figure 2, S4): Mouse anti-V5 (1:1000) (Abcam ab27671) and mouse anti-FLAG (1:1000) (Sigma-Aldrich F1804). To address depleted supply of the original chicken HTP-3 antiserum, new chickens were immunized with a recombinant His6-tagged protein fragment corresponding to the C-terminal 331amino acids of HTP-3, as in MacQueen et al. (2005); affinity-purified antibody exhibiting immunostaining signal/specificity comparable to the original was used at 1:500 dilution.

### Imaging

Images of whole-mount or nuclear spread fixed germ line preparations were acquired on a DeltaVision OMX Blaze microscopy system with a 100 x 1.4 NA widefield objective. Z-stack images (200 nm z-steps) were deconvolved and registration corrected with SoftWoRx software and, where applicable, were stitched together using the “Grid/Collection” plugin on Fiji/ImageJ (Preibisch et al. 2009). For live images of dissected gonads, worms were dissected in 14 µL Egg Buffer containing 0.1% tween-20 and 0.5 mM sodium azide to inhibit movement, and images were acquired using a Leica Stellaris 8 system with a 63x 1.3 NA APO objective and a z-step of 300 nm. 3D-Structured Illumination Microscopy (SIM) images were acquired using a 100x NA 1.40 objective on the OMX system using 3 angles, 5 phases, and a z-step of 125 nm. SIM images were 3D reconstructed and corrected for registration using SoftWoRx. For display, images were maximum-intensity projected to encompass a single layer of whole nuclei. In Fiji, the display pixel minimum/maximum was adjusted for clarity using the Brightness/Contrast tool and images were exported as RGB TIFs for figure assembly. For experiments involving quantification of fluorescence intensity, samples were prepared, imaged, processed, and displayed using the same conditions and settings.

### Quantification of Cytological Features

**Rad51 Focus Quantification** was performed as in Yamaya et al. (2023), with modifications. In brief, Z-stack images of whole gonads were maximum-intensity projected to display a single layer of germ cell nuclei, and these projections were oriented horizontally with the distal tip to the left. Individual non-overlapping nuclei were manually identified beginning in the mitotic tip and ending just prior to diplotene and were marked at the approximate center of the nucleus using the Fiji/ImageJ multipoint selection tool. The XY value of each selected point was used to define the position of the nucleus within the gonad (i.e. the x-axis values in Figure 1I). The selected nuclei were manually segmented via the polygon selection tool to create a region of interest (ROI) and the ImageJ “Find Maxima” program was applied within each selection to generate the number of foci per region. For Find Maxima parameters, the “Prominence” value was determined empirically, starting at the max value of background fluorescence as determined by manual sampling, and then iteratively running the plugin with different ‘Prominence” values to minimize the numbers of false positive and false negative foci as visually identified. All other Find Maxima parameters were maintained as defaults. Two whole mount gonads were quantified for each genotype.

**MSH-5^pT1009^ signal intensity quantification** was performed using a rectangular field spanning the onset of late pachytene (spanning the transition from nuclei with multiple MSH-5 foci to nuclei with 6 prominent MSH-5 foci) for Figure 3, or with rectangular fields from the early, late, and (if available) “later” pachytene regions for Figure 4. Foci in the MSH-5^pT1009^ channel were marked via the Fiji/ImageJ plugin “Find Maxima” and the Peak Intensity (Raw Integrated Density for a single maxima pixel) for each focus was recorded via the Fiji “Measure” function. For Figure 3, peak intensity was recorded for the second channel (*i.e.* COSA-2) using the same marked points as in the MSH-5 ^pT1009^ channel. Background was recorded as the average intensity of 15 randomly selected non-nuclear pixels within the field. For Find Maxima parameters, the “Prominence” value was determined as described above for RAD-51 quantification. At least 3 gonad spreads each were assessed for *cosa-2* mutant and control.

**MSH-5 quantification along meiotic progression** was performed on maximum intensity projected nuclear spread gonad samples containing both early and late pachytene regions. Each stitched image was oriented so that meiotic progression proceeded from left to right; only well-separated nuclei were analyzed. The location of the approximate centroid of each nucleus was recorded, and its x coordinate was used to define its nuclear position along the axis of meiotic progression. For each gonad examined, nuclear positions were normalized relative to the length of the region of the gonad sampled, with the left-most nucleus analyzed designated as the zero position for normalization and the right-most nucleus analyzed corresponding to 100%. Numbers of MSH-5 foci in each nucleus were quantified as described above for RAD-51 foci, and the rolling median number of MSH-5 foci per 10 consecutive nuclei was plotted as a function of relative nucleus position. To account for any differences in the regions of gonads sampled, the midpoint of the X-axis was standardized to be the onset of late pachytene (defined as the first position at which the rolling median of 10 nuclei was 6 MSH-5 foci per nucleus for at least 3 consecutive measurements). 3 gonad spreads each were assessed for *cosa-2* mutant and control.

**Number of MSH-5 foci per chromosome pair** was quantified in nuclei containing 6 MSH-5 foci per nucleus as determined by Find Maxima (see previous section). An ROI encompassing each nucleus was manually segmented and isolated as a 3-dimensional (3D) Z-stack image. Blind to the MSH-5 channel, individual resolvable HTP-3 tracks were manually identified using Z-stack through-scanning, aided by the Fiji/ImageJ 3D Projection plugin (default settings). Individual chromosomes were considered resolvable if a single HTP-3 track could be followed from start to end. Following identification of resolvable HTP-3 tracks, the MSH-5 channel was overlayed and the number of MSH-5 foci on each resolvable chromosome track was recorded. 110 control and 144 *cosa-2* resolvable HTP-3 tracks from 38 nuclei per genotype were scored.

**MSH-5 doublet quantification** was performed similarly to Woglar and Villeneuve (2018). Foci were scored as doublets, elongated foci, or singlets from maximum projection images of spread nuclei from the centers of three SIM fields positioned just after the onset of late pachytene. Foci were scored as doublets if two peak intensities could be visually distinguished while displaying the MSH-5 channel using the “mpl-inferno” LUT in Fiji/ImageJ. Foci were scored as elongated if the focus was clearly oblong, but two separate peaks were not clearly discernable. Rarely, foci signals detected during widefield imaging were no longer detectable with SIM imaging. 147 MSH-5 foci in control and 146 MSH-5 foci in *cosa-2* were scored from 25 nuclei in each condition.

**SUN-1^pS8^/ CHK-2 active zone quantification** was performed on whole mount immunostained germ lines using the 40x air objective on a Zeiss Axio Imager.M2 microscope system. Germ lines were computationally straightened using the “Straighten” function in Fiji/ImageJ on a segmented line manually applied along the approximate midline of the germ line. Using these straightened images, we quantified length of the meiotic SUN-1^pS8^-positive zone (*i.e.* the region of contiguous rows of nuclei in which the majority of nuclei exhibit SUN-1^pS8^ staining) relative to the length of the meiotic prophase zone, defined here as beginning at meiotic prophase entry and ending where the number of nuclei per row decrease to 1-2 (*i.e.* diplotene). At least 10 germlines were quantified per genotype.

### Sample preparation for Western blot experiments or proteomics

Frozen adult worm “popcorn” was prepared following the protocol described in Zanin et al. (2011). Strains were synchronously grown in liquid culture at 22.5°C until the young adult stage. Worms were harvested by sucrose flotation and snap-frozen in liquid nitrogen. The frozen worm pellets were ground into a fine powder using a mixer mill (Retsch). The resulting powder were resuspended in lysis buffer (50 mM HEPES, 160 mM NaCl, 90 mM KCl, 2 mM EDTA, 0.5 mM EGTA, 4 M urea, 0.2% SDS, 0.5% sodium deoxycholate, 1% Triton X-100, 1 mM PMSF, protease inhibitor cocktail tablet [Millipore Sigma, cat. #11873580001], pH 7.5). The suspension was vortexed and sonicated at 30% amplitude using a microtip (Branson, SFX550). Lysates were centrifuged at 15000 rpm (Beckman Coulter, JA-17) for 20 minutes at 4°C, and the supernatant was passed through a 0.22 µm filter. In some experiments, lysates were subjected to overnight dialysis using Thermo Scientific Slide-A-Lyzer Dialysis Cassettes (10K MWCO) to remove free biotin prior to bead binding. To enrich biotinylated proteins, the filtered lysate was incubated with magnetic streptavidin-coated beads (Pierce, cat. #88817) pre-washed with lysis buffer. Protein samples were rotated with beads overnight at 4°C. The beads were subsequently washed twice with 50 mM Tris-HCl (pH 7.5), twice with 2 M urea in 50 mM Tris-HCl, and once with 50 mM Tris-HCl, changing to new microcentrifuge tubes for each wash. Beads were then resuspended in 50 mM Tris-HCl and used immediately for Western blot analysis or stored at −20°C until further processing for mass spectrometry.

### Western Blot

Bead-bound samples were mixed with sample buffer and heated at 95°C for 5 minutes. Proteins were separated on 4-15% precast gradient SDS-PAGE gels (Bio-Rad cat #4561084) and transferred to nitrocellulose membrane using the Trans-Blot Turbo system (Bio-Rad). Membranes were blocked in 5% (w/v) non-fat milk (Bio-Rad, cat. #1706404) in PBST (13.7 mM NaCl, 0.27 mM KCl, 1 mM Na_2_HPO_4_, 0.18 mM KH_2_PO_4_, and 0.1% Tween 20, pH 7.4) for 1 hour at room temperature, followed by overnight incubation with primary antibodies diluted in the same blocking solution. Blots were wash three times in PBST for 10 min each, then incubated with secondary antibodies for 1 hour at room temperature. After three additional 10-minute PBST washes, membranes were developed using Clarity Western ECL Blotting Substrates (Bio-Rad) and imaged.

### Protein Digestion and Mass Spectrometry Analysis

Biotinylated proteins bound to streptavidin on beads were reduced in 5 mM dithiothreitol in 10 mM TEAB for 45 min at 60C, cooled to RT and alkylated in 10 mM iodoacetamide in 10 mM TEAB in the dark at RT for 20 minutes. Beads were washed with 10 mM TEAB. Proteins were digested on-bead with Trypsin/LysC (Pierce) at a ratio of 20:1 in 100uL 25mM TEAB buffer at 37°C overnight. Resulting peptides were separated from the beads and dried using vacuum centrifugation. Dried peptides were resuspended in 2% acetonitrile/0.1% formic acid and analyzed by nanoflow reverse phase chromatography coupled with tandem mass spectrometry (nLCMS/MS) on an Orbitrap Exploris 480 mass spectrometer (Thermo Fisher Scientific) interfaced with a Vantage Neo UHPLC (Thermo Fisher Scientific). Peptides were separated on a 15 cm × 75 μm i.d. self-packed fused silica columns with ReproSIL-Pur-120-C18-AQ (2.4 um, 120 Å, Dr. Maisch, ESI Source Solutions) using an 2-90% acetonitrile gradient over 85 minutes in 0.1% formic acid at 300 nl per min and electrosprayed through a 30 µm stainless steel emitter tip (Bruker) at 20 kV applied post column. Survey scans (MS) of precursor ions were acquired with a 3 second cycle time from 375-1500 m/z at 120,000 resolution at 200 m/z with 1e6 standard automatic gain control (AGC) target and an auto maximum injection time (IT). Precursor ions were individually isolated within 1.6 m/z by data dependent monitoring and 45s dynamic exclusion and fragmented using an HCD activation collision energy 30. Fragmentation spectra (MS/MS) were acquired using a 1e5 Standard AGC and auto maximum injection time (IT) at 30,000 resolution.

### Data analysis

Tandem mass spectra were extracted by Proteome Discoverer (v2.4, ThermoFisher Scientific) and analyzed using Mascot (Matrix Science, London, UK; version 2.8.3). Mascot was set up to search the Caenorhabditis_elegans_20240205 database (UniProt 2024, 26695 entries) assuming the digestion enzyme trypsin. Mascot was searched with a fragment ion mass tolerance of 0.020 Da and a parent ion tolerance of 10.0 PPM. Carbamidomethyl of cysteine was specified in Mascot as a fixed modification. Carbamyl of cysteine and lysine, phospho of serine, threonine and tyrosine, GG of lysine, biotin of lysine and the n-terminus, SUMO2135 of lysine and SUMO3549 of lysine were specified in Mascot as variable modifications. Scaffold (version Scaffold_5.1.2, Proteome Software Inc., Portland, OR) was used to validate MS/MS based peptide and protein identifications. Peptide identifications were accepted if they could be established at greater than 95.0% probability by the Peptide Prophet algorithm (Keller et al. 2002) with Scaffold delta-mass correction. Protein identifications were accepted if they could be established at greater than 95.0% probability and contained at least 1 identified peptide. Protein probabilities were assigned by the Protein Prophet algorithm (Nesvizhskii et al. 2003). Proteins that contained similar peptides and could not be differentiated based on MS/MS analysis alone were grouped to satisfy the principles of parsimony.

### Protein Degradation

Auxin-induced protein degradation (AID) was performed as previously described by (Zhang et al. 2015). Briefly, animals were cultured under standard conditions until transfer to NGM plates containing 2 mM indole-3 acetic acid (IAA, auxin analog) for durations ranging from 30 minutes to 24 hours, as described in each experiment. For auxin experiments involving DAPI quantification or immunofluorescence, animals were harvested at 26 hours post L4. Each experiment included an NGM control and a 24 hour 2 mM auxin plate control to validate effective degradation via presence of expected phenotypes.

### Statistics

Statistics were performed in GraphPad Prism. Details for each statistical analysis are available in the corresponding figure legend.

### Competing Interests

The authors have no competing interests.

## Acknowledgements

We thank C. Akerib and R. Yokoo for genetic screening efforts, C. Akerib for strain constructions, and members of the Villeneuve lab for many fruitful discussions. We thank D. Libuda, A. Dernburg, V. Jantsch and N. Silva for generously sharing antibodies, strains, and/or plasmids. We thank the Caenorhabditis Genetics Center (CGC), funded by NIH Office of Research Infrastructure Programs (P40 OD010440), for strains. We acknowledge K. Lee and the Cell Sciences Imaging Facility (CSIF) for microscopy support; CSIF is supported, in part, by Award Number 1 S10 OD01227601 from the National Center for Research Resources (NCRR) (OMX) and by NIH ORIP funding Award Number 1 S10 OD032300-01 (Stellaris). We also thank R. Cole in the Mass Spectrometry and Proteomics Core at Johns Hopkins School of Medicine for mass spectrometry analysis.

## Author Contributions

CJU – Conceptualization, Investigation, Analysis, Writing (original draft), Writing (review and editing), Funding Acquisition

DYD – Investigation, Analysis, Writing (review and editing)

AMV – Conceptualization, Investigation, Analysis, Writing (original draft), Writing (review and editing), Funding Acquisition

YK – Analysis, Writing (original draft), Writing (review and editing), Funding Acquisition

## Funding Sources

This work was supported by NIH grant R35GM126964 to AMV, NIH grant F32GM153100 to CJU, and NIH grant R35GM124895 to YK.

